# A 36-base hairpin within lncRNA *DRAIC*, which is modulated by alternative splicing, interacts with the IKKα coiled-coil domain and inhibits NF-κB and tumor cell phenotypes

**DOI:** 10.1101/2024.12.23.629241

**Authors:** Xiaoxiao Hao, Yuechuan Chen, Divya Sahu, Róża K. Przanowska, Mujawar Aaiyas, Chase A. Weidmann, Isaac Nardi, Kevin M. Weeks, Anindya Dutta

## Abstract

A tumor-suppressive long noncoding RNA (lncRNA) DRAIC (<u>d</u>own-regulated RN<u>A</u> in <u>c</u>ancers) inhibits NF-κB activity and physically interacts with IKKα, a kinase component of the IKK complex, in several cancer types. Here we explore the precise molecular mechanism involved in this interaction and suppression. Using SHAPE-MaP, we identified a 36-nucleotide hairpin (A+B) within DRAIC that is necessary and sufficient for its anti-oncogenic function. RNA immunoprecipitation (RIP) and Electrophoretic mobility shift assays (EMSA) confirmed this hairpin physically interacts with the coiled coil domain of IKKα. A+B RNA has a high binding affinity (KD ∼1-7 nM) to the coiled-coil domain of IKKα. The binding of A+B disrupts the dimerization of NEMO and IKKα coiled-coil domains, a critical step for IKK action. Consistent with this, A+B inhibits the phosphorylation of the NF-κB inhibitor IκBα and suppresses NF-κB activity. Publicly available tumor RNAseq data revealed that alternative splicing modulates the presence of this critical hairpin: the inclusion of exon 4a (encoding one side of the A+B hairpin) in lung tumors correlates with reduced NF-κB activity. By demonstrating that the A+B hairpin is both necessary and sufficient to inhibit IKK and oncogenic phenotypes, this study underscores the centrality of IKKα interaction and NF-κB inhibition in DRAIC-mediated cancer suppression and indicates that the activity of this lncRNA is regulated by alternative splicing. This study also reveals the first example of a short RNA disrupting coiled-coil dimerization, offering a new approach to disrupt such dimerization in cancer biology.

## Introduction

Long non-coding RNA (lncRNA) are transcripts >200nt in length that do not code for open reading frames >50 amino acids. Increasing evidence has shown that lncRNAs are significant regulators of gene expression, involved in a variety of signaling pathways that drive cancer progression [1, 2]. Unlike protein coding mRNA, the function of a lncRNA relies on the intricate secondary and tertiary structure of the RNA. For example, the 5’ end stem-loop structure within lncRNA XIST is responsible for the recruitment of specific chromatin modifiers [3, 4], while the triple-helix structure at the 3’ end of MALAT1 stabilizes its transcription and is involved in splicing [5, 6]. Similar structural modules are being discovered [7–9] or await discovery [10–15] in many other interesting lncRNAs. Despite the increasing recognition of the importance of lncRNAs in cancer biology, the structural features that underline their functional specificity have largely remained unexplored.

Many lncRNA have multiple isoforms, whether arising from alternative splicing, promoter usage or post-transcription modification. The isoform diversity often leads to the inclusion or exclusion of critical functional domains that substantially alter the function of the lncRNA. For instance, the shorter isoforms of MALAT1 generated by using alternative polyadenylation lose the 3′ triple-helix structure, impairing their ability to localize to nuclear speckles and regulate pre-mRNA splicing [6]. Similarly, there are two existing NEAT1 isoforms: NEAT1_1 (shorter isoform) and NEAT1_2 (longer isoform). The longer NEAT1_2 isoform contains a scaffold that is required for paraspeckle assembly, while the shorter one lacks this region and has different functions [7].

Among the lncRNA involved in cancer progression, the downregulated non-coding RNA in cancers (DRAIC) was first discovered in our lab as a tumor suppressor that inhibits tumor cell migration, invasion and xenograft tumor formation in prostate cancer and glioblastoma cell lines [16–19]. We demonstrated that the same 204 base region of DRAIC exerts its tumor suppressive function and interrupts the IKK complex, which is the key upstream regulator of NF-κB signaling pathway. However, we will not know if the tumor suppressive function is entirely dependent on the interaction with IKK until we map the two activities to the same small structural module [16, 20]. The DRAIC: IKK interaction was remarkably specific, with DRAIC failing to interact in the cell with many equally or more abundant proteins (NF-κB, IκBα, TAK1, TBK1, STAT3 or Ago), and conversely, IKK not interacting with equally or more abundant RNAs (PCGEM, Linc00152, MALAT1, PCA3, PCAT1, SchLp1, GAPDH, or GS1). Thus, defining the smallest functional module of DRAIC will help future studies explore the structural basis of this specificity. Finally, DRAIC expression predicts good outcome in multiple different cancer types [16, 17, 21–23], suggesting that it has potential as a therapeutic RNA, but here again we need to define smaller functional modules of the RNA, because it is easier to deliver small RNAs to tumors.

DRAIC, with significant implications in cancer progression, displayed great isoform diversity. Yet no study has explored the biological differences among the 76 different isoforms of DRAIC. It is important to first understand which small structural modules of DRAIC are required for interaction with IKKα, NF-κB inhibition and suppression of cancer phenotypes, before we can determine which splice isoforms of DRAIC are functional or not. Because of the great diversity of which isoform of DRAIC is dominant in a given tumor, this information will be critical for determining the function of DRAIC in clinical specimens.

To determine the smallest structural module that mediates DRAIC action, we used SHAPE-MaP-based prediction of likely independent structural modules, followed by structure-function analysis to pinpoint the minimal functional region of DRAIC responsible for its interaction with IKK and subsequent inhibition of NF-κB activity. A 36-base sequence within DRAIC, encompassing nucleotides 705-722/741-758 (termed A+B), that can fold into a hairpin is critical for this interaction. This region effectively suppresses cancer cell invasion, migration, and clonogenicity by physically interacting with IKKα and thus inhibiting NF-κB activity. By defining a very small specific interaction region of DRAIC and its target and finding that this same region is critical for inhibiting NF-κB and key cancer pathways, we strongly suggest that NF-κB inhibition is the key pathway by which DRAIC exerts its anti-cancer function. The high affinity (K_D_ <5 nM) of this short hairpin for IKKα positions this part of DRAIC as a potential therapeutic molecule for cancer treatment. This critical region of DRAIC is contained in an alternatively spliced exon, the inclusion or exclusion of which defines which splice isoforms are capable of regulating NF-κB driven gene expression in lung adenocarcinomas. Finally, this is the first demonstration that a short RNA can disrupt dimerization mediated by coiled-coil domains, identifying a novel tool for therapeutically disrupting such interaction in many areas of cell biology.

## Results

### SHAPE-MaP informed modeling of DRAIC RNA structure

To identify functional motifs within the DRAIC RNA, we chemically probed the RNA structure of DRAIC extracted from cells with the SHAPE reagent 5NIA. To identify SHAPE-reactive nucleotides, we employed mutational profiling reverse transcription (MaP-RT), where SHAPE-modified nucleotides trigger skipping or random nucleotide addition in product cDNA, and these “mutation” events can be measured by sequencing. MaP cDNA amplicons generated by SuperScript II RT are on average ∼500 nucleotides in length, so we targeted our initial MaP experiments to the functional part of DRAIC (705-950 bp), ultimately amplifying DRAIC nts 655-1132 and measuring SHAPE reactivity for nts 677-1107. We modeled the RNA structure of DRAIC including our SHAPE reactivity as added restraints [24–26] (**Fig. 1**). We discovered three very well-determined RNA structures within the active region (modeled pairing probabilities > 95 %, **Fig. 1**) that we considered for functional elaboration, including a multi-bulge-containing duplex containing regions we termed A, B, and C (**Fig. 1**). While this ABC region is among the most well-determined in our model, high SHAPE reactivity in stem A suggests that this structure may be somewhat intrinsically flexible, at least in the absence of proteins and other macromolecules.

**Fig. 1.**
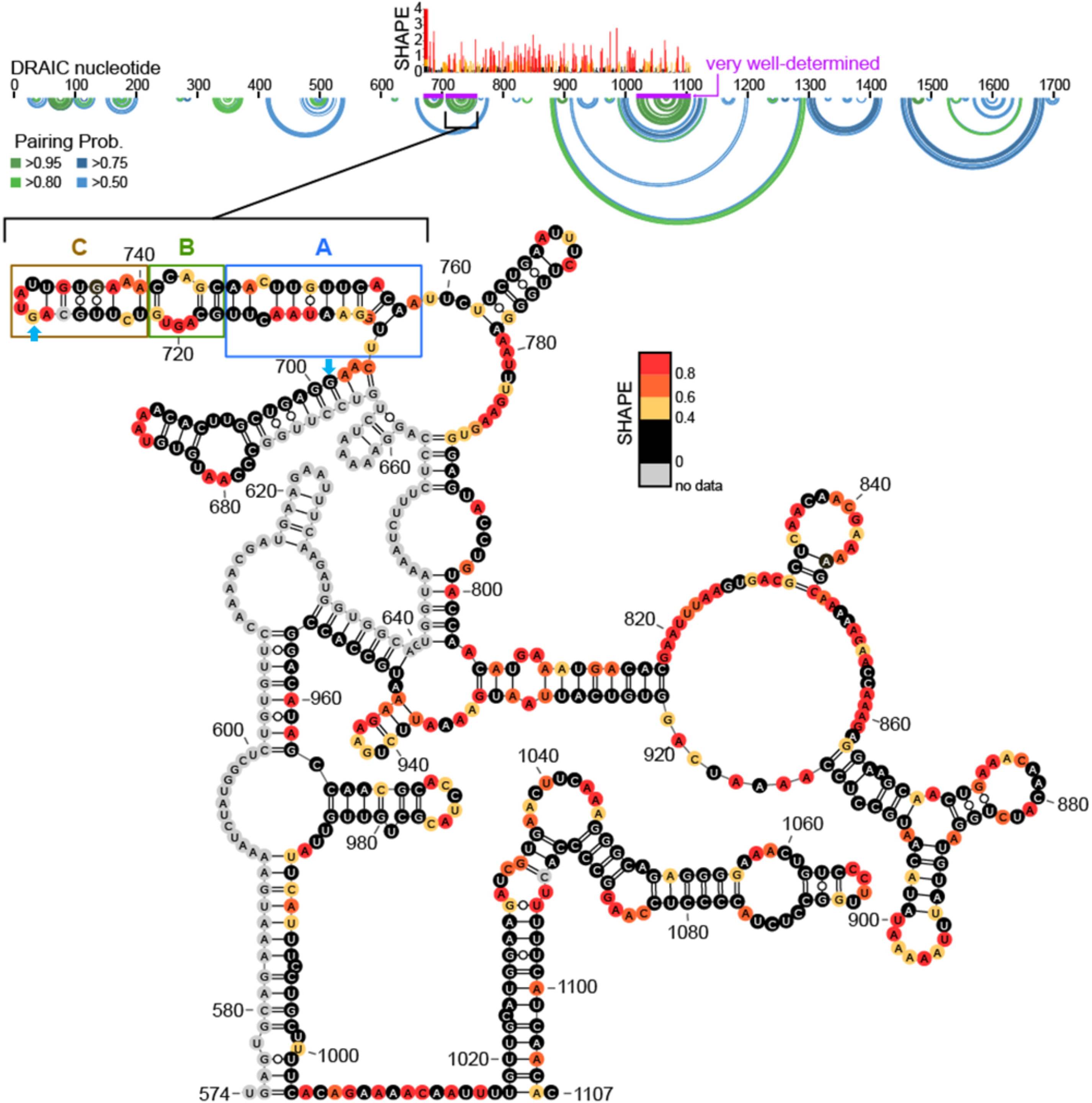
Secondary Structure and SHAPE-MaP Analysis of functional DRAIC. The secondary structure of the functional region of DRAIC, spanning nucleotides 574–1107, is illustrated with SHAPE reactivity data. SHAPE reactivity is shown as a color gradient from black (no data) to red (high reactivity, 0.8), indicating nucleotide flexibility. Notably, the region from 574–676 is computationally predicted to pair with corresponding regions. Regions of interest, including sections A, B and C are highlighted with blue, green, and orange boxes, respectively. Pairing probabilities for various nucleotides are visualized using circular plots, with different levels (>0.50 in light green, >0.75 in dark green, >0.8 in light blue, and >0.95 in dark blue) shown in the corresponding rings. The purple-colored regions indicate well-determined structural areas, which include the locations of the ABC regions, highlighting their structural stability within the DRAIC sequence. The sequence between the blue arrowheads is the exon 4a, that is excluded in some DRAIC splicing isoforms.

### A 36-base hairpin containing nucleotides 705-722/741-758 (A+B) is sufficient and necessary to inhibit NF-κB activity

We dissected DRAIC into several fragments based on the modeled secondary structure and used NF-κB luciferase reporter assays to identify the regions responsible for inhibiting NF-κB activity (**Fig. 2A**). Consistent with our published data, we found that both the full-length (FL) DRAIC and the 705-950 fragment significantly inhibit NF-κB activity compared to the empty vector (EV) and a 900-1705 fragment of DRAIC [16]. When we systematically truncated the 705-950 region of DRAIC, we discovered that the ABC region (nts 705-758) was sufficient for inhibiting NF-κB. Further deletions within the ABC domain revealed that the C portion of the hairpin (nts 722-741) is not necessary for the inhibition because regions A and B expressed as a single fused hairpin maintained full activity. All constructions containing the natural A+B sequence, as well as the fused A+B hairpin alone, showed significant inhibition of NF-κB activity in three different cell lines (**Fig. 2B**).

**Fig. 2.**
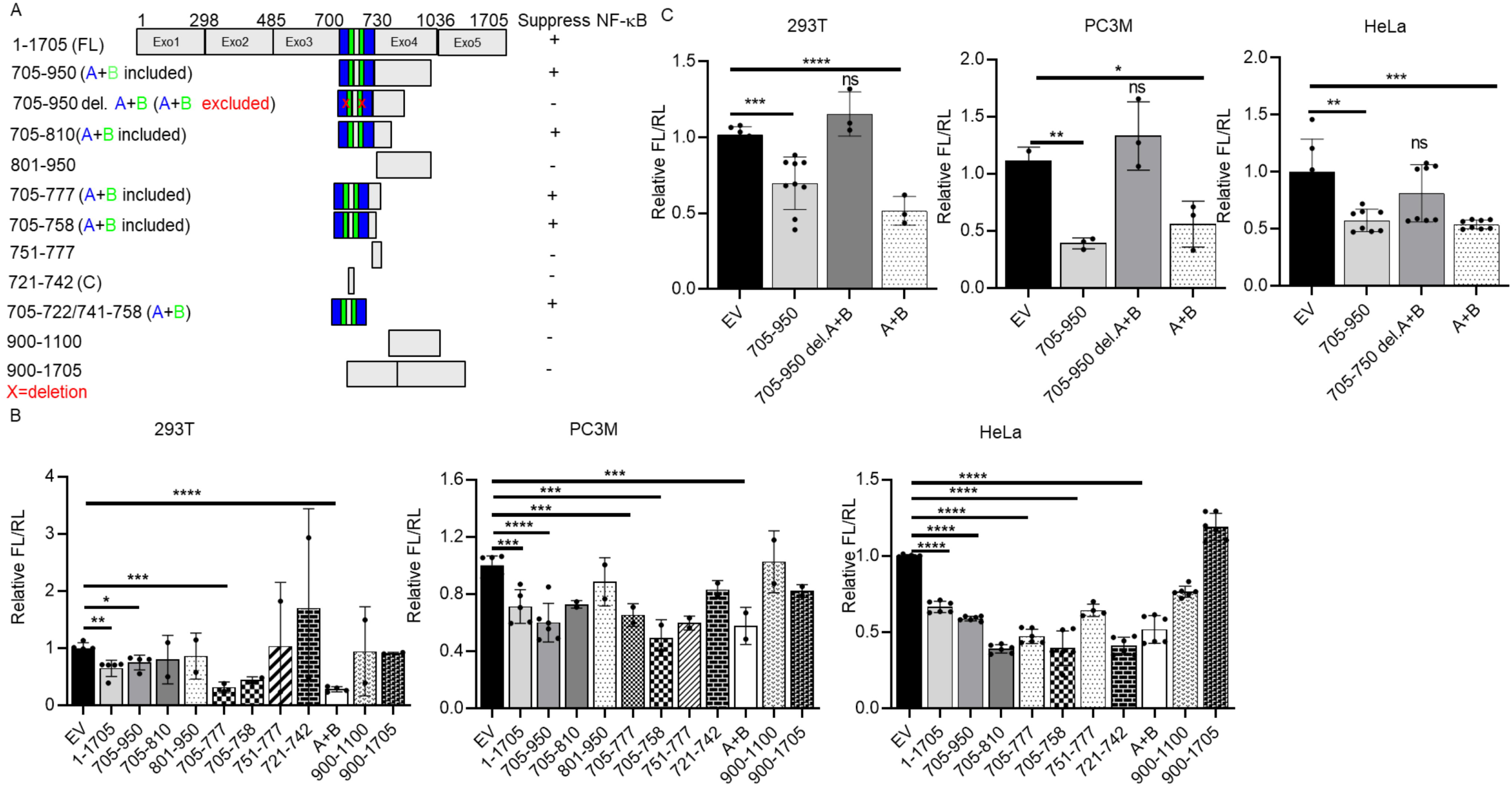
36-bases contained in the A+B region of DRAIC is necessary and sufficient to inhibit NF-κB activity. **A** Schematic of full-length DRAIC and deletion constructs and a summary of their ability to inhibit NF-κB driven luciferase reporter. The blue and green portions constitute A+B. “+”: inhibits NF-κB activity. “-”: does not inhibit NF-κB activity. **B-C** NF-κB driven luciferase reporter assay (Firefly luciferase/Renilla luciferase) in HEK293T, PC3M, and HeLa cells transfected with empty vector (EV) or plasmids overexpressing different portions of DRAIC. Data are expressed as mean ± SD. \**P* < 0.05, \*\**P* < 0.01, \*\*\**P* < 0.001, \*\*\*\**P* < 0.0001 by One-tailed, unpaired Student *t*-tests (B) or by one-way ANOVA followed with Tukey correction (C). ns=no significant. In Figure 2B, only the significant comparisons are marked. Unmarked bars were not significantly different from the EV.

We next tested whether the A+B region was required for NF-κB inhibition (**Fig. 2C**). Deletion of A and B sequence from the functional region of DRAIC (nts 705-950) abolished repression in three cell lines, and we again confirmed that A+B region alone is sufficient to inhibit NF-κB. Together, these results demonstrate that the A+B hairpin is both necessary and sufficient for the suppression of NF-κB activity.

### A+B selectively interacts physically with IKKα, but not NEMO or IKKβ

Next, we explored if this minimal A+B region is necessary for DRAIC RNA to interact with IKKα. We performed RNA Immunoprecipitation (RIP) assays, where in-vitro-transcribed RNA and the bacterially produced recombinant IKKα protein are mixed *in vitro.* The IKKα protein was immunoprecipitated by IKKα antibody or IgG (negative control) followed by RT-qPCR to detect the co-immunoprecipitation of the RNA with the protein (**Fig. 3A**). Luciferase RNA was used as a negative control RNA that does not interact with IKKα. Consistent with previously published data, we observed FL DRAIC and RNA containing nts 705-950 of DRAIC displayed significant enrichment with IKKα. Further dissection of nts 705-950 showed that the 705-758 fragment, containing the prominent ABC hairpin associates with IKKα. Although we observed the 705-777 fragment exhibits higher enrichment to IKKα compared to FL and 705-950 DRAIC, this is not seen in other independent experiments and so we are not sure whether 758-777 has additional sequence that interacts with IKKα. Interestingly, when A+B (the proximal part of the ABC hairpin) was deleted from 705-950 (705-950 del A+B), the ability to bind to IKKα was lost, suggesting that A+B is necessary for interaction with IKKα.

**Fig. 3.**
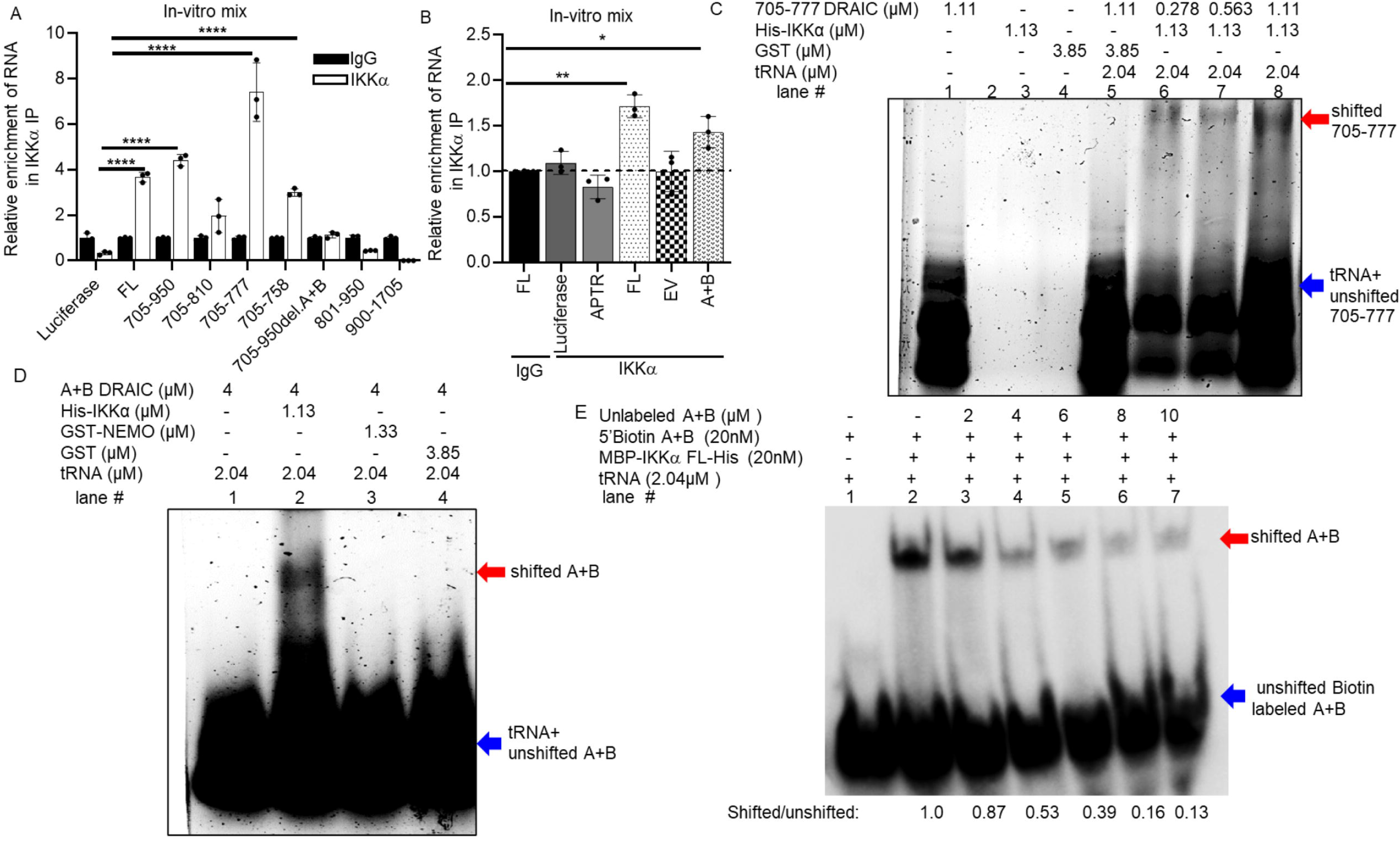
Hairpin structure A+B specifically physically interacts with IKKα *in vitro*. **A-B** In vitro–transcribed (IVT) DRAIC RNA fragments were incubated with recombinant IKKα protein and pulled down with IgG or IKK α antibody followed by RT-PCR of the RNA to see which portion of DRAIC interacts with IKK α protein. Luciferase RNAs, APTR lncRNA, and a short sequence from empty vector (EV) were used as negative controls. The relative enrichments were normalized to IgG. **C-D** Electrophoretic mobility shift assays (EMSAs) to see if DRAIC (705-777) and A+B DRAIC RNAs physically interact with IKKα protein. GST or GST-NEMO were used as negative controls. tRNAs were used as non-specific competitors. The red arrowhead indicates the protein-bound RNA (“shifted” RNA), blue arrowhead indicates the free RNA (“unshifted” RNA). **E** Competitive EMSA was performed to see the specific interaction of A+B DRAIC RNA to IKKα protein. 5’ Biotin labeled A+B DRAIC RNA was incubated with IKKα in the presence of unlabeled excess A+B DRAIC RNA as competitor. The arrowheads are the same as in **C-D**. The numbers at the bottom of the image are the ratio of shifted to unshifted RNAs which is normalized to lane #2. Data are expressed as mean ± SD. \**P* < 0.05, \*\**P* < 0.01, \*\*\**P* < 0.05, \*\*\**P* < 0.0001 by One-tailed, unpaired Student *t*-tests (B and C).

We next tested whether the A+B RNA alone can bind to IKKα (**Fig. 3B**). Here we used APTR lncRNA, luciferase RNA and the cloning site insert RNA from EV (same size as A+B RNA) as negative controls and found that A+B alone can interact with IKKα. Therefore, the A+B part of the hairpin was necessary and sufficient for interaction with IKKα *in vitro*.

To visualize the protein-nucleic acid complex, we employed gel electrophoretic mobility shift assay (EMSA) where the indicated recombinant proteins were mixed with the indicated DRAIC RNA fragment in the presence of a vast excess on non-specific RNA (tRNA) [27, 28]. Total RNA was visualized after electrophoresis of the mix on an agarose gel by SYBR-Green staining. Compared to the no-protein-added lane (lane #1), a shifted band was observed in the presence of IKKα in an RNA dose-dependent manner (lane #6 & 7 & 8) (**Fig. 3C**). No shifted band was observed with GST protein alone (lane # 4), demonstrating some specificity of the 705-777 DRAIC RNA for IKKα. Notably no RNA contamination was observed in GST and IKKα proteins, as there were no signals detected in lane #3 and lane #4.

We next tested whether A+B alone can interact with IKKα by EMSA. A shifted band was observed only in the presence of IKKα (lane #2), but not with GST-tagged NEMO (lane #3) or GST alone (lane #4) (**Fig. 3D**).

The specificity of the interaction was validated by competition experiments. Biotin-labeled A+B RNA probe was used in the EMSA, and the probe was visualized by transfer to membranes and HRP-conjugated streptavidin. A shifted band is seen (lane #2, red arrow), which was competed out by unlabeled A+B duplex in a dose-dependent manner (lane #3-7) (**Fig. 3E**). Concurrently, there was an increase in free biotin-labeled RNA running in the lower part of the membrane (blue arrow). The ratio of shifted to unshifted A+B RNA (normalized to the ratio in lane #2) is shown at the bottom of the image.

We also examined the specificity of the interaction by incubating A+B RNA with 10-fold excess of other recombinant proteins (MBP-Spot, MBP-ZUP1-Spot, and GST-IκBα1-54) instead of IKKα. A+B is bound specifically by IKKα, but not by MBP-Spot, MBP-ZUP1-Spot or GST-IκBα1-54 (**Sup. Fig. S1A**). Conversely, we found that IKKα did not interact with M3 RNA which is the same size as A+B and contains a stem-loop structure but has a different sequence (**Sup. Fig. S1B** and **the M3 RNA sequence provided in the Supp. Table 1**).

**Table 1.**
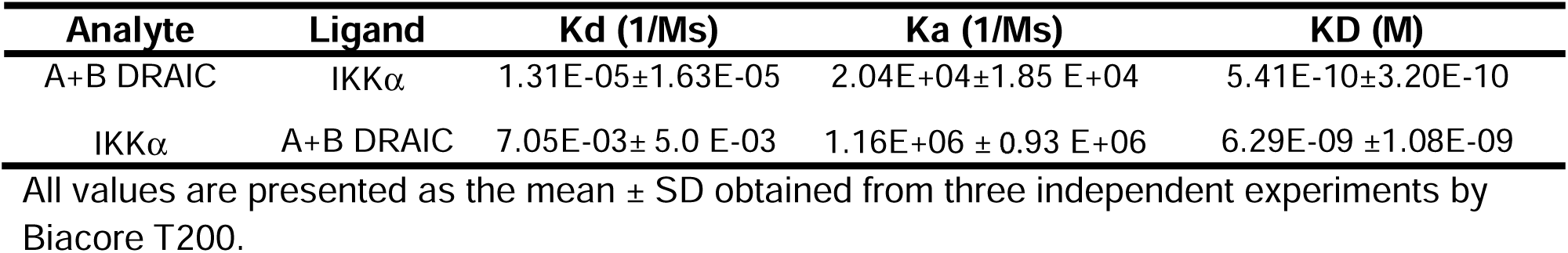
The binding affinity parameters of A+B RNA and IKKα measured by Biacore T200.

| Analyte | Ligand | Kd (1/Ms) | Ka (1/Ms) | KD (M) |
| --- | --- | --- | --- | --- |
| A+B DRAIC | IKKα | 1.31E-05±1.63E-05 | 2.04E+04±1.85 E+04 | 5.41E-10±3.20E-10 |
| IKKα | A+B DRAIC | 7.05E-03± 5.0 E-03 | 1.16E+06 ± 0.93 E+06 | 6.29E-09 ±1.08E-09 |

All values are presented as the mean ± SD obtained from three independent experiments by Biacore T200.

These results demonstrate that A+B RNA specifically binds to IKKα protein and not other proteins, and IKKα specifically binds to A+B RNA and not other RNAs.

### A+B is necessary and sufficient for interaction with IKKα coiled-coil domain, but not IKKβ coiled-coil domain

Our published data showed that DRAIC moderately decreases the association of IKKα with NEMO [16]. The binding of the regulatory and substrate-interaction subunit NEMO of the IKK complex with kinase subunits IKKα or IKKβ is mediated by interactions between <u>c</u>oiled-<u>c</u>oil <u>d</u>omains [20, 29].Therefore, we wondered whether the inter-subunit interaction domain of the IKKα subunit, specifically the coiled-coil domain, interacts with DRAIC.

We first tested if the IKKα <u>c</u>oiled-<u>c</u>oil <u>d</u>omain (IKKα-ccd) binds to DRAIC. A RIP assay after *in vitro* incubation of the RNA and the protein showed that both FL DRAIC and the 705-950 fragment of DRAIC were enriched in the immunoprecipitates of the MBP-IKKα-ccd (**Fig. 4A**). However, no enrichment was observed with either of the negative control RNAs luciferase or APTR RNAs (**Fig. 4A**). We next tested whether hairpin A+B is necessary for DRAIC binding to the IKKα-ccd. Removing hairpin A+B from 705-950 abolished the enrichment of 705-950 DRAIC with MBP-IKKα-ccd. No RNA enrichment was observed with the MBP tag alone or with MBP-IKKα-ccd incubated with negative control RNAs, APTR, or luciferase (**Fig. 4B**). These results demonstrate that the binding of DRAIC to IKKα-ccd requires the presence of hairpin A+B.

**Fig. 4.**
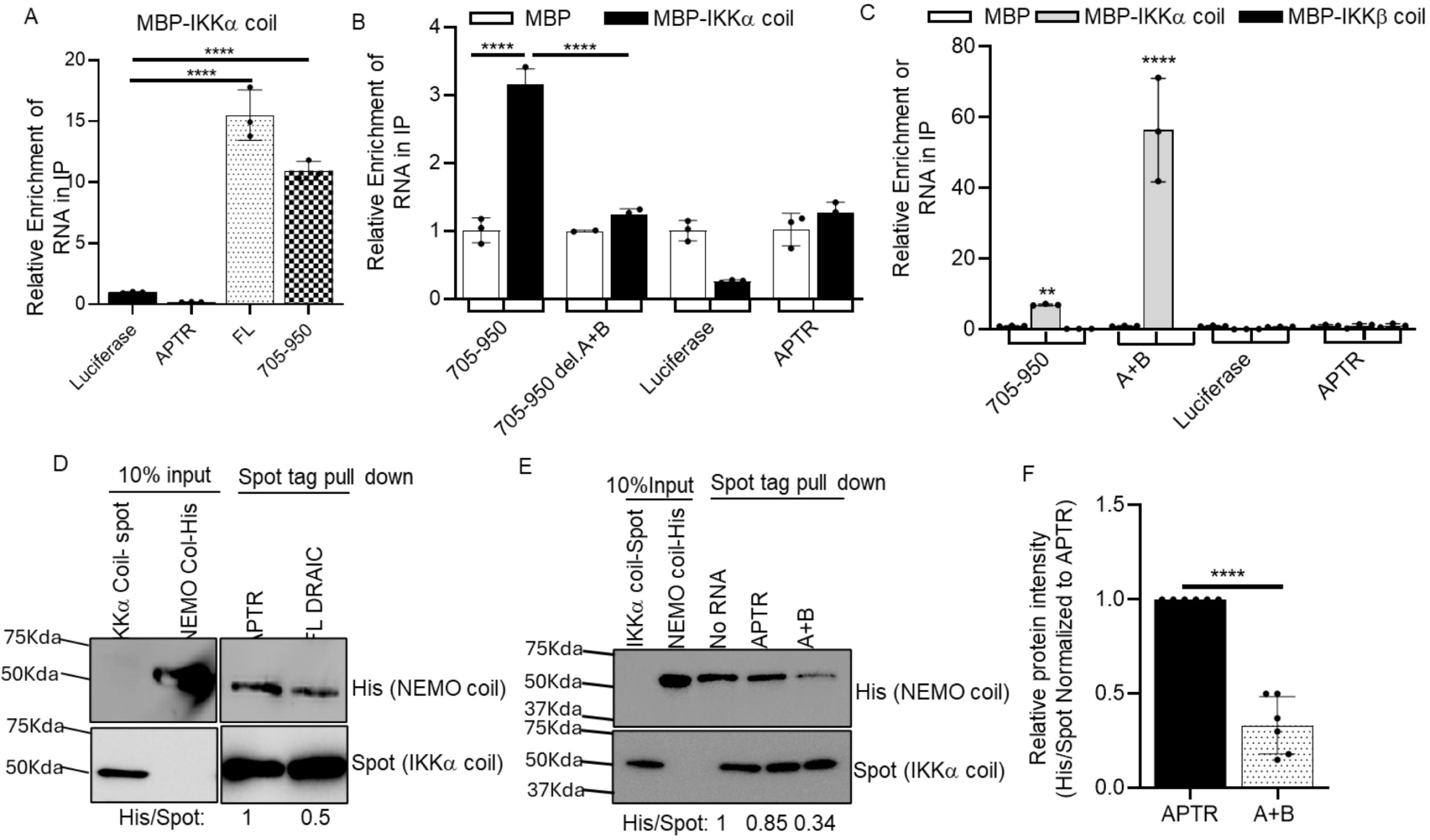
Hairpin structure A+B specifically physically interacts with IKKα coiled-coil domain. **A-C** RIP to detect if FL DRAIC and indicated parts interact with IKKα coiled-coil domain. Luciferase and APTR lncRNA were used as negative controls. MBP tag only and MBP-IKKβ coiled-coil domain protein were also included as negative controls. **D-E** In vitro IKK complex formation assay to see if the binding of FL DRAIC (D) and A+B DRAIC (E) to IKKα coiled-coil domain affects the binding of NEMO to IKKα. IKKα coiled-coil protein was immobilized on Spot magnetic beads, then FL DRAIC RNA or A+B DRAIC RNA and His-NEMO coiled-coil protein were introduced. Proteins associated with the Spot beads are visualized by Western blotting. **F** The quantification of the relative protein intensity of Spot (IKKα coiled-coil) and His (NEMO coiled-coil) in the presence of APTR and A+B DRAIC RNA. Data are expressed as mean ± SD of 6. \*\**P* < 0.01, \*\*\**P* < 0.05, \*\*\*\**P* < 0.0001 by one-way ANOVA followed with Tukey correction (A), two-way ANOVA followed with Tukey correction (B and C) or by one-tailed, unpaired Student *t*-tests (F).

Additionally, we produced MBP-IKKβ-ccd protein, containing the same number of amino acids as the IKKα-ccd, and carried out RIP assay. DRAIC 705-950 was associated specifically with IKKα-ccd but not the IKKβ-ccd, (**Fig. 4C**). Interestingly, hairpin A+B alone displayed a high level of enrichment specifically with the MBP-IKKα-ccd, but not the MBP-IKKβ-ccd or MBP tag alone. These results demonstrate the specific binding of DRAIC 705-950 with IKKα-ccd, and that A+B hairpin alone is sufficient for this specific interaction.

### A+B decreases the binding of NEMO coiled-coil to IKKα coiled-coil domain

Because A+B interacts with the coiled-coil domain of IKKα, we investigated whether the binding of hairpin A+B to the IKKα-ccd inhibits the binding of the NEMO-ccd to IKKα-ccd.

MBP-IKKα-ccd-Spot protein was immobilized on Spot beads and incubated with MBP-NEMO-ccd-His6 protein in the presence of FL DRAIC or negative control RNA, APTR. A significant decrease in NEMO-ccd pulled down by IKKα-ccd was observed in the presence of FL DRAIC RNA compared to APTR RNA (**Fig. 4D**, left). The hairpin A+B was then added instead of FL DRAIC, and this was sufficient to inhibit the interaction between the two IKK proteins (**Fig. 4E, F**). Therefore, the A+B RNA is sufficient to inhibit the interaction between the coiled-coil domains. Since the A+B RNA does not interact with the GST-NEMO (**Fig. 3D**) we believe that the inhibition is carried out through the interaction of the RNA with IKKα-ccd.

### A+B binds to IKKα with high affinity and inhibits the phosphorylation of IκBα

Surface Plasmon resonance [30, 31] was used to measure the affinity of IKKα with A+B RNA. His6 tagged IKKα was immobilized and A+B RNA injected through the flow cell as the analyte. A dose-responsive binding was seen, and the calculated K_D_ of the interaction was 0.54±0.32 nM (**Fig. 5A, and Table 1**). When the reverse experiment was done with 5’biotin-tagged A+B bound to the sensor surface as the ligand and IKKα injected as the analyte, we still saw dose responsive interaction, but the calculated K_D_ was 6.09±1.08 nM (**Fig. 5B, and Table 1**). The lower affinity when the RNA is tethered to the chip could be because of the absence of a spacer between the chip surface and the A+B RNA, leading to steric hindrance of the association with IKK protein [32]. Note that the bulk-shift that occurs at the end of the association phase in Fig. 5B, when the analyte buffer is replaced by the running (washing) buffer does not affect the measurement of the K_D_ because the curve-fitting model is based on the curves during the active association and dissociation stages. Together the results suggest that the K_D_ of the interaction is in the range of 1-5 nM. SPR confirmed that A+B RNA does not interact with the NEMO (**Fig.** 5C).

**Fig. 5.**
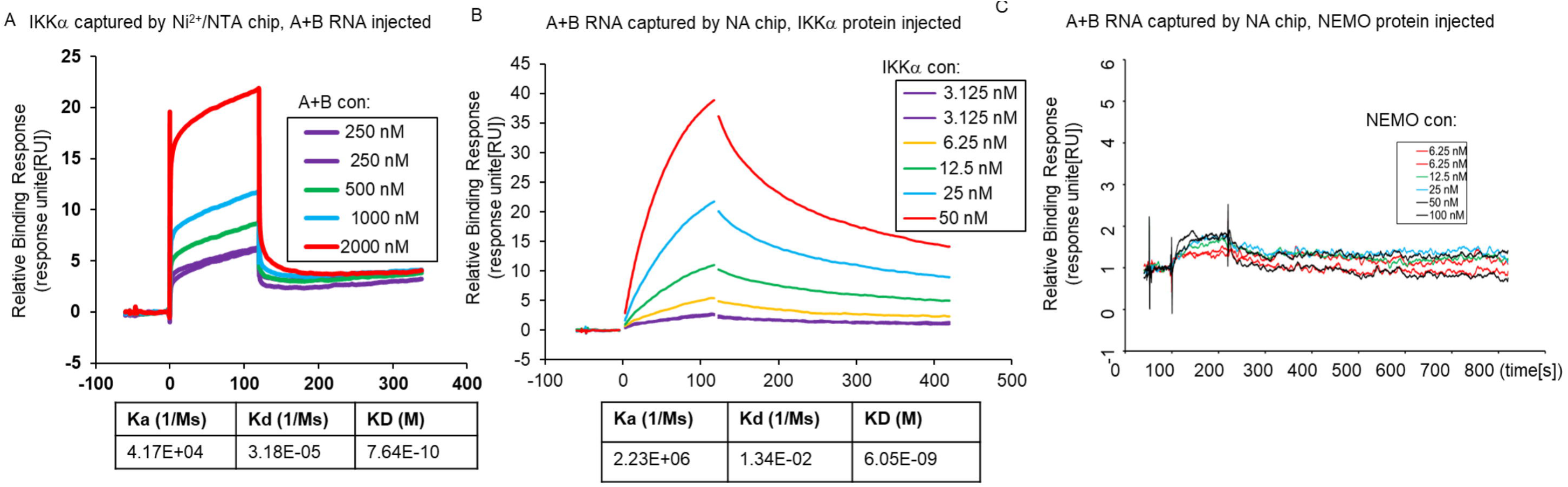
High binding affinity of A+B to IKKα measured by Surface Plasmon resonance. **A** His6-tagged IKKα was immobilized on the sensor chip using Ni^2+^/NTA chip, and doses of A+B RNAs injected. The table at the bottom of the sensorgram shows the kinetic parameters that are on-rate K_a_ and an off-rate K_d_ and the calculated K_D_ (K_d/_ K_a_) based on 1:1 steady binding model. **B** 5’ Biotin-A+B RNA was immobilized on the sensor surface by NA (Neutravidin) chip, and doses of MBP-IKK α-His protein were injected in solution as the analyte. All the experiments were conducted at 25°C. **C** Similar to B, doses of GST-NEMO protein were injected over the sensor chip after 5’Biotin-A+B RNA was immobilized.

### A+B binds to IKKα, disrupts the IKK complex, and inhibits the phosphorylation of IκBα when expressed in cells

We performed a RIP assay in 293T cells after overexpressing (OE) different fragments of DRAIC, using IKKα immunoprecipitation (IP) from cell extracts to assess whether DRAIC or fragments thereof are co-precipitated with IKKα. Immunoprecipitation of the same lysate with IgG was used as a negative control, and the amount of each RNA present in the IKKa IP was normalized to the amount in the IgG IP. Western blotting showed that equal amount of IKKα was precipitated in each lane (**Fig. 6A**). There was significant enrichment of FL, 705-950 and A+B RNA in the IKKα precipitates compared to the IgG negative control (**Fig. 6B**). Moreover, the A+B element is necessary (compare 705-950 with and without A+B) and sufficient (A+B alone) to interact with IKKα *in vivo* (**Fig. 6B**).

**Fig. 6.**
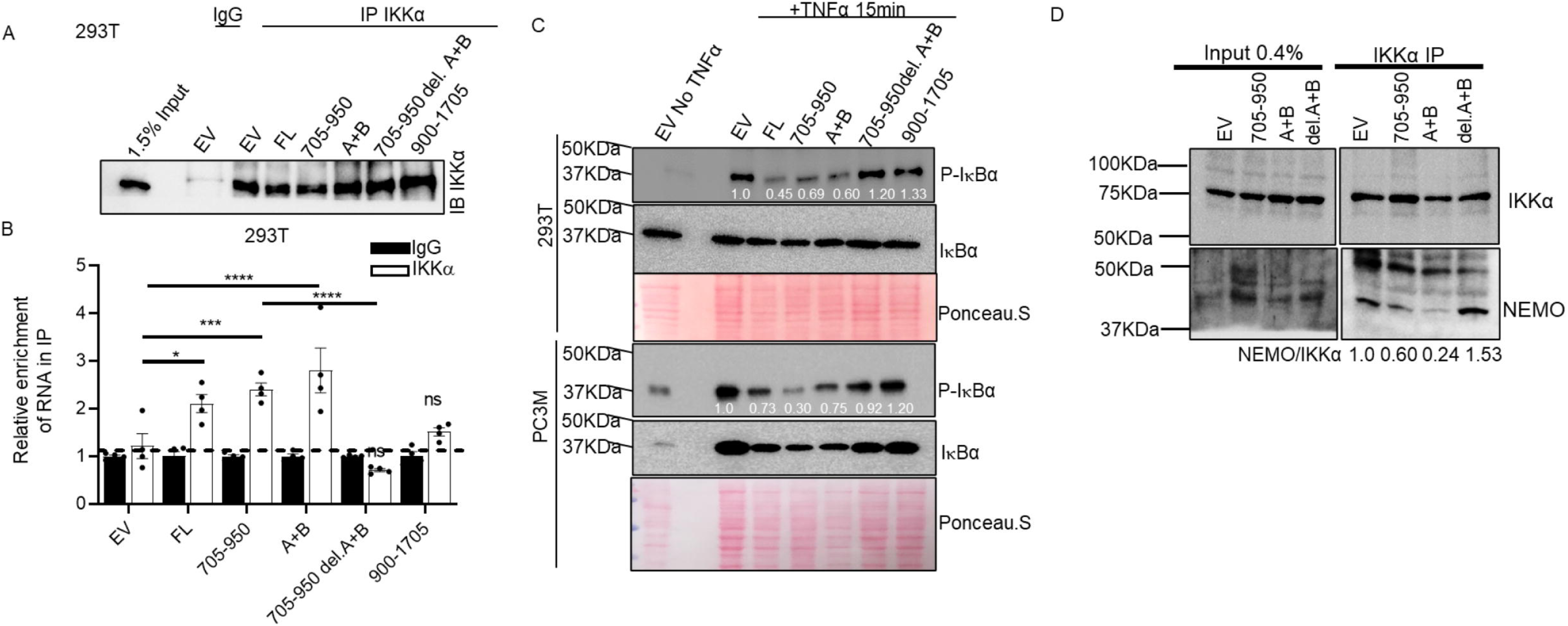
A+B binds to IKKα and inhibits the phosphorylation of IκBα, and disrupts the interaction between NEMO and IKKα when expressed in cells. **A-B** RIP was performed after overexpression of FL DRAIC and DRAIC derivatives to see if A+B part of DRAIC interacting with IKKα in vivo. (A) Immunoblotting of IKKα in the immunoprecipitates showing equal pull down of the protein. (B) RT-qPCR showing the relative enrichment to IKKα (normalized to IgG). **C** Western blotting was performed to detect the phosphorylation levels of IκBα after overexpressing DRAIC derivatives in HEK293T (upper panel) and PC3M (lower panel). Ponceau S. staining was used to adjust the equal amount of protein loaded. The ratio of p-IκBα to total IκBα, after normalization to the ratio in the EV lanes is displayed below the blot. **D** To assess whether the A+B fragment of DRAIC disrupts the interaction between NEMO and IKKα, we overexpressed different DRAIC fragments and performed immunoprecipitation of IKKα followed by detection of NEMO in the immunoprecipitated samples. The NEMO:IKKα ratio in the immunoprecipitates is a measure of the association between the two proteins.

Next, we tested which derivative of DRAIC can inhibit the kinase activity of IKK in cells by measuring the phosphorylated level of IκBα. Although the total level of IκBα did not change (because its degradation was prevented by MG132), the addition of TNFα stimulated its phosphorylation [33]. We quantitated the phospho-IκBα relative to total IκBα to show that the phosphorylation was inhibited by FL DRAIC and 705-950 DRAIC **(Fig. 6C).** Deletion of A+B from 705-950 DRAIC (705-950 del. A+B) prevented the inhibition, while the A+B RNA alone was sufficient to inhibit the phosphorylation. Similar results were seen in four different cell lines suggesting the universality of these observations (**Fig. 6C, Supp. Fig. S2A and 2B**). Thus, the A+B RNA is necessary and sufficient to inhibit the kinase activity of IKK *in vivo*, consistent with A+B being necessary and sufficient to physically associate with IKKα.

In our previously published data [16], we showed the DRAIC disrupt the binding of NEMO to IKKα in vivo. Consistent with this, we also observed that 705-950 and A+B DRAIC significantly decreased the co-immunoprecipitation of NEMO with IKKα (**Fig. 6D**). However, removal of A+B from 705-950, inhibited the ability to disrupt the NEMO: IKKα interaction.

### A+B inhibits cancer cell migration, invasion, and clonogenicity

We next investigated whether A+B affects tumor cell behavior similarly to FL DRAIC. We transfected various DRAIC fragments (FL, 705-950, 705-950 del. A+B, A+B only, and 900-1705) into several cancer cell lines (HeLa, PC3M, and C4-2B) and measured the impact of A+B on tumor cell behavior. RT-qPCR confirmed the successful overexpression of different DRAIC fragments in cells (**Supp. Fig. S3, Fig. S4B and S4D**).

In the Matrigel Invasion Assay, the tumor cells are seeded in a chamber containing a Matrigel extracellular matrix and asked to invade through the matrix to the bottom surface of the membrane[34]. The images of the cells and the quantitation of the results are shown (**Fig. 7A-B and Supp. Fig. S4Aand S4C**). Consistent with our previously published data, FL and 705-950 DRAIC significantly inhibited tumor cell invasion[16]. The A+B DRAIC fragment alone significantly decreased tumor cell invasion and removing A+B from 705-950 completely abolished this effect, demonstrating that A+B is sufficient and necessary for DRAIC to inhibit tumor cell invasion (**Fig. 7A and Supp. Fig. S4Aand S4C**). This pattern was observed in three different cell lines.

**Fig. 7.**
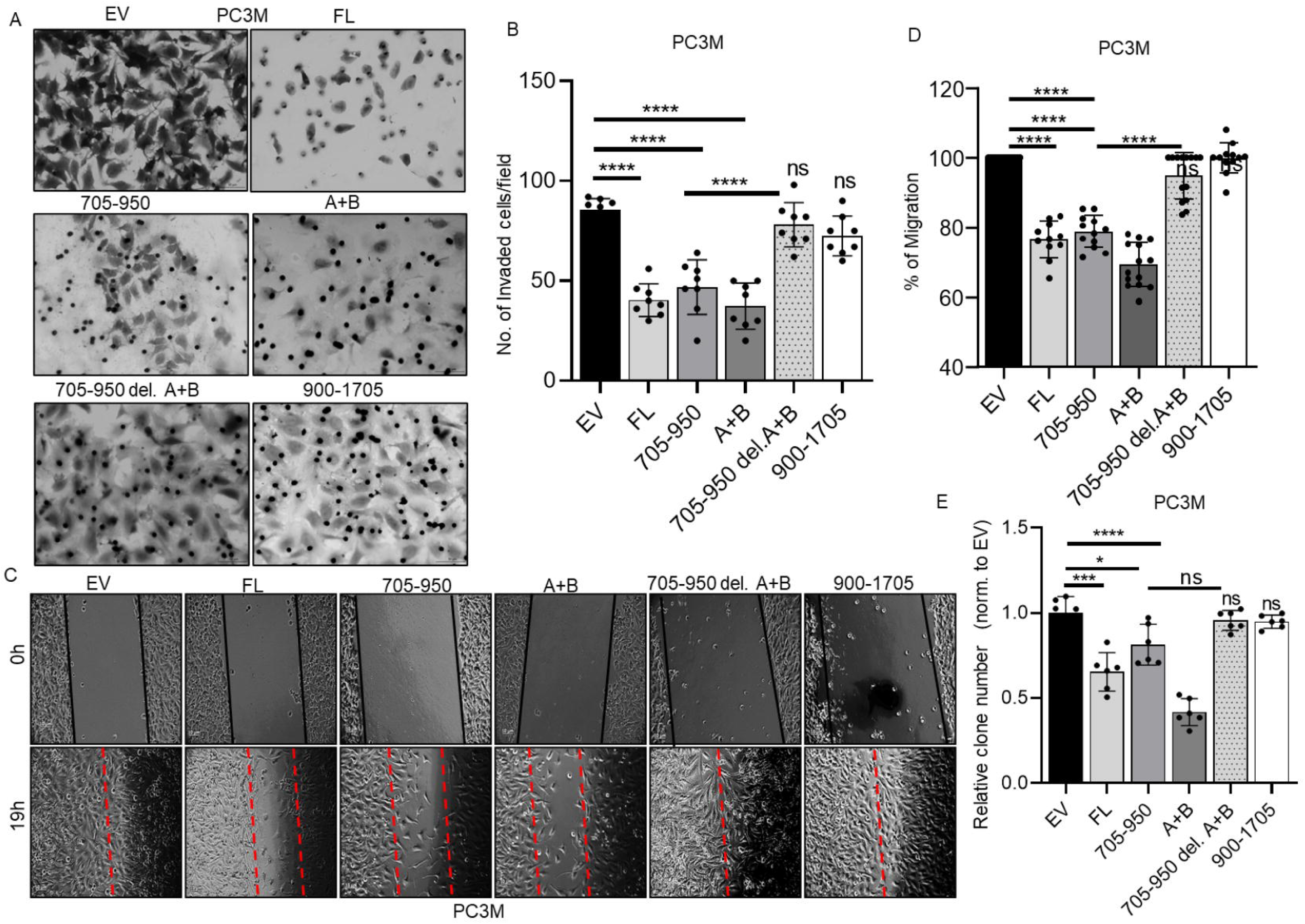
A+B is required for DRAIC to suppress cancer cells migration, invasion and clonogenicity when expressed in cells. **A** Matrigel Invasion assay was conducted in PC3M cells after transfection with FL DRAIC and DRAIC derivatives to see if A+B affects tumor cell invasion. The scale bar is 50 μm. **B** Quantitation of the invaded cells from A. **C** Wound Healing assay was performed to evaluate if A+B affects PC3M cell migration after DRAIC transfection. The dotted lines define the areas lacking cells. The scale bar is 10 μm. **D** Quantitation of the wound closure from C, normalized to the closure with EV. **E** Quantitation of the colony number after DRAIC in Clonogenicity assay to see if A+B affects PC3M cell clonogenic growth on plastic tissue culture plate. Cells were fixed and stained with 0.5% Crystal Violet. Data are expressed as mean ± SD of 2 replicate; 4-5 random fields captured for each replicate; \**P* < 0.05, \*\*\**P* < 0.001, \*\*\*\**P* < 0.0001 by one-way or two-way ANOVA followed with Tukey correction. ns= not significant when compared to EV.

Next, we tested if A+B affects tumor cell migration by performing a wound-healing assay[35]. As displayed in **Fig. 7C** and quantitated in **Fig. 7D**, the gap (“wound”) was almost closed in both EV and 900-1705 DRAIC, while significant gaps remained in FL, 705-950, and A+B DRAIC. Removing A+B from 705-950 nullified the inhibition of migration by 705-950.

Lastly, we tested if A+B influences the clonogenic cell growth in culture. Tumor cells were seeded on tissue culture plates, and we observed that FL, 705-950, and A+B significantly decreased the number of colonies compared to EV and 900-1705 DRAIC **(Fig. 7E).** However, removing A+B from 705-950 abolished this inhibition.

Taking all these results together, the A+B part of DRAIC is necessary and sufficient to inhibit tumor cell invasion, migration, and clonogenic growth.

### A+B is contained in an alternative exon 4a, whose inclusion or exclusion differs between cell lines

An examination of RNAseq data from cell lines revealed an alternative splicing event with an alternate splice acceptor site that adds a 5’ extension (exon 4a) to the constitutive exon 4 (**Fig. 8A**). Interestingly, exon 4a encodes the bottom strand of the A+B+C hairpin (**Fig. 1**).

**Fig. 8.**
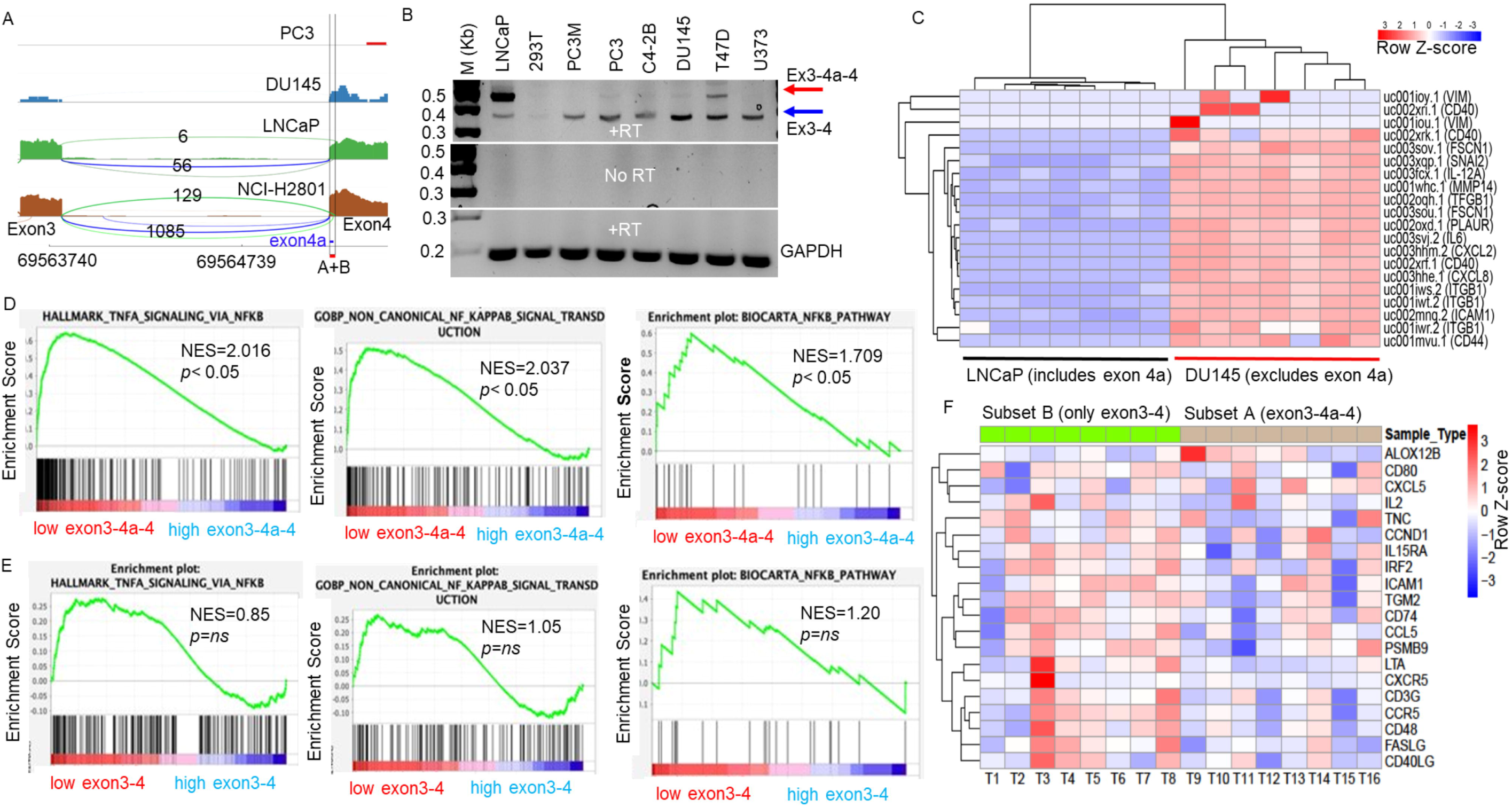
Alternative splicing regulates the ability of DRAIC to inhibit NF-κB signaling in cancer cell lines and in LUAD tumor. **A** The sashimi plot of RNA-seq data illustrates the alternative splicing event involving exon 4a in DRAIC across various cell lines. The lines connecting exon 3 and exon 4 represent splicing events, either directly between exon 3 and exon 4 (green line) or through the inclusion of exon 4a (blue line). The numbers along the lines indicate the frequency of these splicing events. The red-colored region highlights the A+B segment, with the blue-colored region specifically marking exon 4a. The numbers at the bottom denote the chromosomal loci on chromosome 15. **B** RT-PCR analysis showing the relative abundance of DRAIC transcripts with and without Exon 4a in several cell lines. RT=reverse transcription. M stands for DNA marker. **C** Heatmap showing downregulation of NF-κB-regulated inflammatory genes in LNCaP cells expressing exon 4a compared to DU145 cells lacking exon 4a. The scale represents the Row Z-score. **D** GSEA highlighting NF-κB-regulated inflammatory genes enriched in LUAD tumors in TCGA with low exon 3-4a-4 expression (Q1) compared to those with high exon 3-4a-4 expression (Q4). NES=Normalized enrichment score. P<0.05 is statistically significant. **E** GSEA shows that NF-κB-regulated inflammatory enriched genes are not differentially enriched in LUAD tumors with low (Q1) or high (Q4) expression of exon3-4 isoform of DRAIC. **F** The heatmap illustrates NF-κB target gene expression in eight LUAD tumors matched by expression of DRAIC but expressing or not expressing exon 4a (from Table 1). NF-κB targets are upregulated in tumors expressing exon 3-4 compared to those expressing exon 3-4a. The scale represents the Row Z-score.

To evaluate the clinical significance of the A+B containing splice isoform, which contains exon 3-4a-4, we analyzed its expression across multiple cell lines. Sashimi plots showed the exon 3-4a-4 DRAIC isoform was predominantly expressed in LNCaP and LUAD NCI-H2801 cells, while PC3 and DU145 cells mainly expressed isoforms lacking exon 4a (designated as exon 3-4 isoforms). RT-PCR confirmed the differential expression of isoforms with and without exon 4a in several cell lines (**Fig. 8B**). Notably no genomic DNA contamination was observed in the cDNA, as there was no signal detected in the RNA samples without RT **(Fig. 8B** the middle panel). RNA-Seq analysis demonstrated reduced expression of NF-κB regulated inflammatory genes in LNCaP (expressing exon 4a) compared to DU145 (not expressing exon 4a) (**Fig. 8C**). Similar trends are also observed in other cancer cell lines (**Supp. Fig. S5**). This is consistent with the hypothesis that DRAIC isoforms that contain exon 4a, resulting in an intact A+B hairpin, can inhibit NF-κB.

### Tumors expressing exon 4a have lower expression of NF-κB target genes

We next turned to tumors showing differential expression of the exon 4a containing isoform. Lung adenocarcinomas (LUAD) were the only tumor type with sufficient tumors expressing either splice junction exon 3-4a-4 or exon 3-4. GSEA was conducted after stratifying tumors based on expression of the exon 3-4a junction or the 3-4 junction. The results show that tumors with lower expression of 3-4a junction had higher activity of NF-κB driven genes compared to tumors with high expression of 3-4a junction (**Fig. 8D**) with Normalized enrichment scores (NES) of 1.7-2.0 (p<0.05). These enrichments were not seen and not significant when the tumors were stratified by low and high expression of 3-4 junction (**Fig. 8E**).

To extend this result, we identified LUADs that exclusively express exon 3-4a junction (with no exon 3-4 junction) (**Subset A in Table 2A**) and vice versa. Only 8 tumors completely skipped exon 4a (**Subset B in Table 2A**) and we selected 8 matching tumors that included exon 4a from Subset A. The tumors were matched by the level of expression of the total DRAIC transcript (**Table 2B**) to eliminate any complications from differential expression of all DRAIC isoforms. A heatmap of NF-κB target genes confirms that although the matched tumors express equal levels of all DRAIC isoforms, expression of exon 4a (Subset A) is associated with decreased NF-κB target gene expression (**Fig. 8F**).

**Table 2:** A) LUAD tumors from TCGA classified by expression of 3-4a-4 junction vs 3-4 junction B) Eight matched tumors from the two subsets matched by equal expression of DRAIC, used for the heatmap of Fig. 7C

|  | **exon3-4a-4** | **exon3-4** | **Total samples** |
| --- | --- | --- | --- |
| Subset A | yes | no | 183 |
| Subset B | no | yes | 8 |

B. DRAIC expression in CPM (Count per million)
|  | **Subset B (exon3-4)**<br>**Tumors 1-8** | **Subset A (exon 3-4a-4)**<br>**Tumors 9-16** |
| --- | --- | --- |
| Tumor 1,9 | 0.38 | 0.38 |
| Tumor 2,10 | 3.92 | 3.92 |
| Tumor 3,11 | 0.83 | 0.82 |
| Tumor 4,12 | 3.83 | 3.91 |
| Tumor 5,13 | 0.69 | 0.67 |
| Tumor 6,14 | 0.56 | 0.55 |
| Tumor 7,15 | 0.36 | 0.36 |
| Tumor 8,16 | 0.41 | 0.41 |

## Discussion

### Identification and Specificity of the A+B RNA Structure

We utilized SHAPE-MaP to model the secondary structure of a key functional region of DRAIC. The DRAIC lncRNA regulates the NF-κB signaling pathway by interacting with the two subunits of the IKK complex. Our findings identified a critical 36-nucleotide hairpin within DRAIC, plays a key role in modulating NF-κB activity, as it is both necessary and sufficient for inhibiting NF-κB signaling by interacting specifically with IKKα. Other studies [36] have identified broad functional or protein binding regions within lncRNAs, generally spanning over 200 nucleotides in length, making discovery of this 36nt A+B hairpin - comparable in size to microRNAs - remarkable. This compact, highly functional region highlights the potential therapeutic significance of identifying minimal structural elements of the RNA for therapeutic purposes. Future studies will focus on determining whether the functionality of this hairpin arises from specific nucleotide interactions or the structural integrity of the hairpin. Notably, other hairpins predicted by SHAPE-MaP at the 3’ end of DRAIC did not exhibit inhibitory activity, indicating the unique specificity of the A+B structure in NF-κB inhibition.

### The interaction of A+B with a specific subunit of IKK

The physical interaction between the A+B hairpin and IKKα, but not NEMO or IKKβ, again supports the specificity of this RNA-protein interaction. This specificity is further corroborated by our findings that A+B directly interacts with the coiled-coil domain of IKKα, a region known to be crucial for the assembly and activation of the IKK complex [20, 37]. Coiled-coil domains, characterized by their intertwined α-helices, are notoriously difficult to disrupt due to their stable hydrophobic core and specific interhelical interactions [38]. These features make the coiled-coil domain of IKKα an especially challenging target for therapeutic intervention. Despite this stability, A+B could efficiently disrupt the binding of NEMO-coiled coil domain to IKKα-coiled coil domains, offering a promising avenue for therapeutic intervention in a wide range of diseases driven by the NF-κB pathway. Many cancers exhibit constitutive activation of the NF-κB pathway, leading to uncontrolled cell proliferation, resistance to apoptosis, and enhanced metastatic potential [39]. By inhibiting the NEMO-IKKα interaction, A+B could reduce NF-κB-mediated transcription of oncogenes and survival factors, making it a potential therapeutic agent in cancers where NF-κB plays a pivotal role. Additionally, A+B could enhance the sensitivity of cancer cells to chemotherapy or radiation therapy by lowering their resistance mechanisms mediated through NF-κB. Clearly, detailed structural studies are needed to understand how A+B disrupts the NEMO-IKKα interaction, e.g. by X-Ray Crystallography.

The inability of A+B to bind to the coiled-coil domain of IKKβ (**Fig. 4C**) supports our previous observation that FL DRAIC RNA bound much better to IKKα than to IKKβ *in vitro*, and disrupted the association of IKKα with NEMO, but not that of IKKβ with NEMO [16]. This selectivity for the α subunit over the β subunit, both of which are catalytic, and both of which could homodimerize with NEMO to form a functional IKK, suggests that one way cancer cells may escape the inhibition of NF-κB by DRAIC is by expressing more IKKβ than IKKα [40, 41] or in rare cases, by activating NF-κB by pathways independent of IKKα [42]. This model could be a potential explanation for why there are some cancers where DRAIC is not tumor suppressive, e.g. breast cancer [43].

NEMO plays a pivotal role in the canonical NF-κB signaling pathway, acting as a regulatory subunit of the IKK complex. The binding of NEMO to IKKα and IKKβ is required for the kinase activation, thus leading to the phosphorylation and activation of NF-κB [44]. In addition, it is also reported that NEMO inhibits the receptor interacting with protein-1 (RIP1) from activating CASPASE-8, thereby preventing apoptosis independent of the NF-κB signaling pathway [45, 46]. Targeting NEMO and NF-κB signaling pathway is a promising strategy for cancer treatment. We have demonstrated that FL DRAIC and DRAIC 701-905 also interacted with NEMO [16]. However, the A+B region of DRAIC does not associate with NEMO (**Fig. 3D and 5C**), implying that there are adjoining parts of the RNA, contained in 701-905 but not A+B, which interact with NEMO. Identifying and characterizing these interaction sites could provide deeper insights into the mechanism by which DRAIC inhibits IKK activation.

### Affinity and mechanism of interaction

Surface plasmon resonance (SPR) has been widely utilized to study the binding affinity of proteins, peptides, and small molecules [30], but its application to the study of lncRNAs and their interactions with proteins is still relatively novel. SPR demonstrated that the binding affinity of A+B DRAIC lncRNA with IKKα is in the low nanomolar range (< 6nM). No binding is detected with GST-NEMO, suggesting that A+B’s interaction is specific to IKKα and the A+B RNA itself. The fast association (higher K_a_) and slow disassociation (lower K_d_) suggest that DRAIC can quickly engage with IKKα and remain bound long enough to effectively disrupt the interaction between IKKα and NEMO, leading to sustained inhibition of NF-κB signaling. Further structural studies are necessary to determine whether this affinity is solely due to interactions with the coiled-coil domain or if other regions of IKKα are also involved. The nanomolar K_D_ for A+B binding to IKKα is comparable to other RNA-protein interactions. For example, Nelson et al. reported a binding affinity of 14 nM for the interaction between MHV N protein and RNA [47], while Chen et al. determined a K_D_ of 2.82 nM for the binding of IBV N protein to the IBV leader [48]. The slight discrepancy between the two K_D_ values in our experiments may be attributed to: (i) the method of ligand immobilization, which could result in improper orientation and reduced accessibility of the binding site, and (ii) the differing surface chemistries of the NTA and NA chips, which may influence their interaction with analytes [49]. RNA-protein interactions can be mediated by the RNA backbone, where the secondary structure plays a critical role, or by specific interactions between nucleotides and amino acids, making the RNA sequence important for complex formation. Future experiments will therefore explore whether the hairpin structure of A+B or specific sequences within A+B are crucial for its high-affinity interaction with IKKα.

### Short RNA as a disruptor of coiled-coil domain interactions

To the best of our knowledge, this is the first instance of a short RNA interacting with such high affinity with a coiled-coil domain and disrupting the interaction of two coiled-coil domains. As has been mentioned above, dimerization mediated by coiled-coil domains is widespread in biology and there have been repeated attempts to discover agents that can disrupt such interaction for therapeutic purposes. Thus, we are excited by the possibility that short RNAs, like A+B, can be designed that will selectively disrupt many other coiled-coil domain interactions for therapy.

### Alternative splicing modulates the activity of DRAIC

To the best of our knowledge this is the first example where alternative splicing of a lncRNA removes a critical functional module of the RNA. The results from the cancer cell lines and the lung adenocarcinomas in TCGA suggest that the inclusion of Exon 4a, which constitutes half of the A+B hairpin, is associated with repression of NF-κB, as is expected from our prior biochemical and cell biological results. An interesting possibility is that physiological or pathological regulation of this alternative splicing event will activate or inactivate a regulator of NF-κB signaling. The results also imply that when assessing the role of some lncRNAs in tumor progression, it is important to assess which splice isoform is expressed, and that some contradictory effects ascribed to a lncRNA in the Literature could stem from different splice-isoforms being expressed. However, LncRNAs often have many splice isoforms, making it difficult to assess which isoforms should be studied separately. Thus, it is important to do structure-function analyses as done here to determine which functional modules are likely to be regulated by alternative splicing and then separately study the functionality of the different isoforms of the lncRNA.

### Inhibition of IKK and Therapeutic Application

The phosphorylation and subsequent degradation of IκBα is a crucial step in the activation of NF-κB, allowing the transcription factor to translocate from the cytoplasm to the nucleus and initiate gene expression [46, 50]. In our study, we found that the A+B hairpin effectively inhibits the phosphorylation of IκBα in cells, thereby blocking a key step in the canonical NF-κB activation pathway. This inhibition likely contributes to the observed decrease in NF-κB activity in cells with high DRAIC expression. If this inhibitory effect can be replicated in vitro using soluble IKK, it would enable us to elucidate the precise mechanism of action—whether the interaction with the coiled-coil domain and the disruption of the IKKα complex is the primary or sole means by which A+B exerts its kinase inhibition.

Given the ability of the A+B hairpin of DRAIC to suppress cancer cell migration, invasion, and clonogenicity, its biological significance is clear. These findings indicate that the A+B hairpin not only plays a crucial role in modulating NF-κB signaling but also serves as a potent inhibitor of aggressive cancer phenotypes. This dual function highlights the therapeutic potential of targeting the A+B hairpin structure in cancers characterized by hyperactive NF-κB signaling. A therapeutic strategy centered on the A+B hairpin could involve designing small molecules, antisense oligonucleotides, or RNA mimics that specifically stabilize or enhance the hairpin’s interaction with IKKα. By doing so, this approach could inhibit NF-κB activation and consequently reduce tumor progression, invasion, and metastasis. Additionally, given the specificity of the A+B hairpin for IKKα over IKKβ and NEMO, targeting this hairpin could provide a more precise therapeutic approach with fewer off-target effects, particularly in cancers where IKKα driven NF-κB signaling is a driving factor.

Moreover, combining this strategy with other therapies that target different components of the NF-κB pathway or other oncogenic pathways could enhance the overall efficacy of cancer treatment, potentially overcoming resistance mechanisms that are common in aggressive cancers. Therefore, the A+B hairpin within DRAIC represents a promising therapeutic agent, offering new avenues for the development of cancer treatments aimed at curbing the malignancy associated with hyperactive NF-κB signaling.

### Study Limitations

Despite the promising findings shown in this study, there are some limitations. First, while it is amazing that we identified a splice isoform that naturally inactivates the A+B hairpin we have not been able to rigorously test the functional importance of splice isoforms with or without this hairpin, due to the paucity of LUAD tumors that completely skip exon 4a. We also have not evaluated whether the inclusion or exclusion of exon 4a in DRAIC changes in response to physiological or pathological stimuli. In our current study, we only focused on the interaction of DRAIC with IKKα, and there could be other interesting proteins that interact with DRAIC (or even A+B) to contribute to the tumor-suppressive function of DRAIC. We have discovered that the anti-invasion, -migration, -clonogenicity function of DRAIC and the interaction with and inhibition of IKK map to the same small part of the RNA, strongly suggesting that the anti-cancer cell functions are mediated by IKK inhibition. However, it is possible that future point mutations in the A+B area may separate the inhibition of IKK from the anti-cancer functions.

Furthermore, SHAPE-MaP provides valuable information for structure prediction of RNA but may not be able to capture all the functional structures in different conditions, especially if the RNA folding changes dynamically in time. There is a chance that additional RNA folding states exist depending on the cellular condition. All the anti-cancer effects of the RNA were measured in cell lines *in vitro*, limiting the assessment of biological evaluation. Future studies in xenografts are required to evaluate these finding in the specific tumor microenvironment. Finally, NF-κB has a critical role not only in cancer biology but also in immunology, and we have not addressed whether DRAIC, A+B or splice isoforms of DRAIC play an important role in the regulation of inflammation and the immune response.

In conclusion, our study identifies a critical secondary structure within DRAIC that specifically interacts with IKKα to inhibit NF-κB signaling and suppress cancer cell aggressiveness. The specificity of the A+B hairpin in binding to IKKα, coupled with its functional effects on NF-κB signaling and cancer cell behavior, highlights the potential of targeting this RNA structure for therapeutic intervention in NF-κB driven cancers. Identifying the A+B functional module also allowed us to identify functionally different isoforms of DRAIC generated by alternative splicing. Future studies should explore the detailed molecular mechanisms by which the A+B hairpin modulates IKKα activity and the broader implications of these findings for cancer therapy.

## Material and Methods

### Cell culture and transfection

Cancer cell lines LNCaP, C4-2B, DU145, PC3, T47D, U373 and PC3M were cultured in RPMI 1640 medium supplemented with 10% FBS, 1% penicillin/streptomycin, 1 mmol/L sodium pyruvate, and 10 mmol/L HEPES buffer. HeLa and HEK293T cells were cultured in DMEM medium supplemented with 10% FBS and 1% penicillin/streptomycin. All the cells were maintained in a humidified cell culture incubator at 37°C in the presence of 5% CO_2_. For NF-κB luciferase reporter assays, cells were co-transfected with NF-κB firefly luciferase reporter plasmid, Renilla luciferase plasmid, and DRAIC fragments using Lipofectamine 3000 (Life Technology, cat. no. 3000001) following the manufacturer’s instructions. Twenty-four hours post-transfection, cells were washed and lysed in 1x passive lysis buffer (Dual-Luciferase Reporter Assay, Promega, cat. no. E1980), and luminescence signals were captured by GloMax 96 microplate luminometer (Promega, cat.no. E6521). For overexpression of different portions of DRAIC in cells, DRAIC genefragments were first cloned into a pcDNA3 backbone as previously described[16]. After sequence confirmation, the plasmids were transfected into cells using Lipofectamine 3000.

### Quantitative Reverse Transcription PCR (RT-qPCR) and Western Blotting

To confirm the successful overexpression of DRAIC fragments in cells after transfection, total RNA was extracted using Direct-zol RNA Miniprep kit (ZYMO Research, cat. no. R2052). cDNA was synthesized from equal amount of RNA by using the PrimeScript RT Reagent kit (Takara, cat. no. RR037). RT-qPCR was performed using an ABI Prism 7500 fast system (Applied Biosystems) with the 2x qPCR Master Mix (NEB, cat. no. M3003). For each gene, the relative expression levels were normalized to glyceraldehyde 3-phosphate dehydrogenase (GAPDH). All experiments were conducted at least 2-3 times in triplicate, with results presented as mean values ± S.E.M. PCR was conducted with primers designed to flank exon 3 and exon 4 to detect isoforms either containing or lacking exon 4a. The reaction was carried out for a total of 28 cycles. All primer information is listed in **Table S1**.

### Phosphorylation of IκBα

For detecting the level of p-IκBα, an equal number of cells was plated in 6-well plate for 10-12 hours. The next day, cells were pretreated with MG132 for 2 hours and then treated with TNFα (20ng/ml) for 15 minutes. After 15 minutes, cells were washed with cold PBS, then 300ul 2XSDS sample buffer (containing 5% β-ME) was added to lyse the cells. Lysates were loaded onto 12% SDS-PAGE gel for protein separation and immunoblotted with anti-phosphor IκBα antibody (Abcam, cat.no. ab133462, 1:4000 dilution). The membrane was transferred by Semi-dry transfer system (Bio-Rad) and followed by Ponceau S staining to see the overall protein loading in each group. The total levels of IκBα was also probed as loading controls using anti-IκBα (CST, cat. No.9242). All gel images were obtained on ChemiDoc^TM^ touch system with Image Lab Touch software (Bio-Rad).

### Clonogenic Assay, Wound Healing Assay, and Matrigel Invasion Assay

To evaluate the ability of a single cell to form a colony, a plastic clonogenic assay was conducted. DRAIC fragment-containing cells were diluted to 500 cells/ml and seeded into a 6-well plate, then continuously cultured for 9 days (with fresh media replaced 5 days after seeding). Colonies were fixed with glutaraldehyde (5.0% v/v), stained with crystal violet (0.5% w/v), and images were acquired by ChemiDoc^TM^, the number of colonies in each well were counted.

To evaluate cell migration, cells were seeded on a 24-well plate at 2.5 × 10^5/well. The next day, after cells reached 100% confluency, wounds were generated using a 10 μL micropipette tip. Floating cells were removed and washed with PBS three times, and complete cell culture media were added to each well. Images (3-5 random fields/well, quadruplicate per group) were captured immediately after media replacement (T = 0) and 20 hours (T = 20) after wound generation at 10× magnification (Leica Microscope). After exporting images, wound areas were measured using ImageJ (NIH). Briefly, the polygon selection tool was used to indicate the wound area, and the areas were quantified (Analysis > Measure). The extent of closure at T20 was calculated by subtracting the area at T20 from the area at T0, and the percentage closure was determined by normalizing the difference to the area at T0 [35, 51].

To evaluate cell invasion, the Matrigel-containing Boyden chamber (Corning, cat.no. 354480) was first rehydrated with serum-free medium at 37°C for 2 hours before seeding the cells. Cells were starved for 8-10 hours, and then a total of 1 × 10^5 cells were seeded in serum-free medium in the top of the chamber. The bottom of the chamber contained full growth medium (10% FBS) as a chemoattractant, and the chamber was incubated at 37°C in the presence of 5% CO_2_. After 19-20 hours, the chambers were gently washed with 1x PBS. The non-invading cells from the upper surface of the chamber were gently removed using Kimwipes or cotton swabs. The invaded cells were treated with 4% PFA for 10 minutes, followed by 0.5% crystal violet staining at 22°C for 15-30 minutes to visualize. The invaded cells were captured under a microscope (Leica or KENYEN) as detailed in the Figure legends.

### RNA immunoprecipitation (RIP), RNA pull-down, Co-Immunoprecipitation (Co-IP) assay

The full-length DRAIC (FL), 705-950, 705-950 del. A+B, 900-1705, and other fragments of DRAIC were prepared by PCR (primers provided in **Table S1**) and used as templates for *in vitro* transcription (IVT) reaction by the T7 promoter using T7 polymerase and the HiScribe T7 High Yield RNA Synthesis kit (NEB, cat. no. E2040L). Template DNAs were removed by treating with DNase I at 37°C for 30 minutes and the IVT RNAs were further purified by RNA purification kit (NEB, cat. no. T2040L). The size of each IVT RNA was also confirmed by agarose gel electrophoresis. To elucidate the binding portion of DRAIC on IKKα (Life Technology cat. no. PR7619B), an RNA pull-down assay was performed as previously described[16]. Briefly, IVT DRAIC RNAs (1 µg/reaction) were heated at 95°C for 2 minutes, chilled on ice for 2 minutes, and then slowly cooled to RT for 15 minutes in RNA folding buffer. Refolded RNA was incubated with His-IKKα (100 ng/reaction) at RT for 1 hour in RNA binding buffer (25 mM HEPES, pH 7.5, 150 mM NaCl, 0.05% NP-40, 10% glycerol, protease, and RNase inhibitors) and RNA-IKKα complex was pulled down by Ni^2+^-NTA Agarose Beads (Qiagen cat. no. 30230) or IKKα antibody. The unbound RNA was removed by wash buffer. RNA was extracted from the complex using TRIzol RNA isolation reagent (Life Technology, cat. no. 15596260). cDNA was synthesized and RT-qPCR was performed to see if DRAIC RNA was enriched in IKKα or not.

To check if A+B DRAIC was interacting with IKKα in cells, 293T cells expressing different portions of DRAIC were lysed in PEB (polysome extraction buffer: 20 mM Tris-HCl pH 7.5, 100 mM KCl, 5 mM MgCl2, 0.5% NP-50) cell lysis buffer freshly supplemented with protease and RNase inhibitors. An equal amount of protein (0.5 mg/reaction) was first pre-cleared with protein A/G agarose beads (Santa Cruz, cat. no. sc2003), followed by incubation with anti-IKKα antibody (CST, cat. no. 2682S, 1:200 dilution) and IgG as a negative control at 4°C for 16-18 hours. The next day, protein A/G agarose beads were added to each tube to immunoprecipitate the related protein. After five washes with NT2 buffer (20 mM Tris-HCl pH 7.4, 150 mM NaCl, 1 mM MgCl_2_, 0.05% NP-40) with protease and RNase inhibitors (NEB, cat. no. M0314S), half of each sample was directly added with 2x SDS sample buffer for immunoblotting with anti-IKKα antibody. The other half of each sample was used to extract RNA using TRIzol reagent following the manufacturer’s instructions. cDNA was synthesized and RT-qPCR was performed using DRAIC-specific primers to see if DRAIC RNA was enriched to IKKα or not.

### *In Vitro* IKK complex formation assay

To evaluate if FL and A+B DRAIC RNA will interfere with the binding of NEMO to IKKα, we conducted the IKK complex formation assay. Briefly, Spot-tagged IKKα coiled-coil domain protein was immobilized on Spot-Trap magnetic beads (Proteintech cat. no. ETDMA). His-tagged NEMO coiled-coil domain and either FL DRAIC or A+B RNA was added and incubated for 2 hours at 4°C. The NEMO associated with IKKα was then pulled down by magnetic beads, and after five washes with NT2 buffer, 2x SDS sample buffer (β-ME freshly added) was added to the complex. The presence of IKKα coiled-coil protein and NEMO coiled-coil protein was detected using anti-Spot antibody (Proteintech cat. no. 26A5, 1:4000 dilution) and anti-His tag antibody (Qiagen, cat. no. 34650, 1:4000 dilution), respectively.

### Cell-free SHAPE-MaP

The general strategy has been described previously [52]. LNCaP cells grew to 80% confluency in two 15-cm dishes. Both plates were washed once in PBS before scraping and lysis in 2.5 mL of proteinase K buffer (40 mM Tris, pH 8, 200 mM NaCl, 1.5% sodium dodecyl sulfate, and 0.5 mg/mL proteinase K). Proteins were digested for 45 min at 23 °C with intermittent mixing. Nucleic acids were extracted twice with one volume of phenol: chloroform:isoamyl alcohol (25:24:1) that was pre-equilibrated with 1.1x RNA folding buffer (110 mM HEPES, pH 8, 110 mM NaCl, 5.5 mM MgCl2). Excess phenol was removed through two extractions with one volume of chloroform. The final aqueous layer was exchanged into 1.1x RNA folding buffer using PD-10 desalting columns (GE Healthcare Life Sciences). The resulting RNA solution was incubated at 37 °C for 20 minutes before being split into two equal volumes. The SHAPE reagent, 5-nitroisatoic anhydride (5NIA, AstaTech) in DMSO was added to a final concentration of 62.5mM to one aliquot, and DMSO added to the other. Samples were incubated at 37 °C for 10 minutes. RNA was precipitated with 1/10 volume of 2 M ammonuim acetate and one volume of isopropanol. After washing with 75% (v/v) ethanol, the resulting pellet was dried and resuspended in 88 mL water. 10 mL of 10x TURBO DNase buffer and four units of TURBO DNase (Thermo Fisher) were added and, the mixture was incubated at 37 °C for 1h. RNA was purified (GeneJET RNA Cleanup and Concentration Micro Kit, Fisher) and eluted into 20 mL of nuclease-free water. 1µg of each RNA sample was subjected to MaP reverse transcription (MaP-RT) [32] using a DRAIC-specific primer (**Table S1**). For second-strand cDNA synthesis, the output DNA was used as a template for PCR with primers made to amplify a region of DRAIC containing the functional fragment, with the product including nt 655-1132. Purified PCR products were measured (Qubit dsDNA HS Assay Kit), and 2 ng were used as a template for 10 additional cycles of PCR to add Illumina-specific sequences and sample barcodes. Size distributions and purities of *DRAIC* amplicon libraries were verified (2100 Bioanalyzer, Agilent). Libraries (about 120 amol of each) were sequenced on a MiSeq instrument (Illumina) using a 250×250 paired end sequencing kit. ShapeMapper2 software was used to counts mutations and generate SHAPE profiles [53], and the SuperFold [54] was used with experimental SHAPE data to inform RNA structure modeling. Secondary structure projection images were generated using VARNA [55].

### Electrophoretic Mobility Shift Assay (EMSA)

To visualize the protein-nucleotide complex, we conducted a gel electrophoretic mobility shift assay (EMSA) using the LightShift™ Chemiluminescent RNA EMSA Kit (Life Technology, cat. no. 20158) according to the manufacturer’s instructions. Briefly, the in vitro transcribed unlabeled 705-777 DRAIC or A+B DRAIC RNA was incubated with IKKα-His or GST-NEMO protein for 25 minutes at 22°C. After 25 minutes, 6x loading dye was added to the reaction mixture, which was then loaded onto a 1.2% agarose gel in 0.5x TB buffer (45 mM Tris, 45 mM boric acid) and run at 90-100 volts for 45 minutes. After electrophoresis, gels were stained for 20 minutes with 2× SYBR Gold (Invitrogen) while shaking. To reduce background signal, the agarose gel was destained in ∼150 mL 0.5× TB for 15 minutes with shaking[27].

Competitive EMSA was performed using biotin-labeled A+B DRAIC RNA (produced by Gene Scripts) in the presence of excess unlabeled A+B DRAIC RNA (homemade). The reaction mixture was loaded onto a 6% DNA retardation gel (Life Technology, cat. no. EC6365BOX) in 0.25X TBE buffer and run at 90-100 volts for 45 minutes. After electrophoresis, the binding reaction was transferred to a positively charged nylon membrane (Life Technology, cat.no. AM10100) by wet transfer in 0.25X TBE buffer at 300 mA on ice for 50 minutes. The membrane was removed from the transfer system and cross-linked at 120 mJ/cm² using a commercial UV-light crosslinker. The membrane was then blocked with the Nucleic Acid Detection Blocking Buffer at 22°C for 15 minutes with gentle shaking. After blocking, the membrane was incubated with the conjugate/blocking solution (1:300 dilution) at 22°C for 15 minutes with gentle shaking. Following 4-5 washes with 1x wash buffer, the membrane was developed using X-ray film or ChemiDoc Image developer with enhanced ECL.

### Surface Plasmon Resonance (SPR)

Surface Plasmon Resonance (SPR) experiments were performed using a Biacore T200 (GE Healthcare) at 25 °C, with HBS-EP serving as the running buffer (0.01 M HEPES, pH 7.4, 0.15 M NaCl, 3 mM EDTA, 0.05% vol/vol Surfactant P20 [GE]). An NTA (nitrilotriacetic acid) chip (Cytiva, cat. No. BR100407), used to immobilize the IKKα-His protein, was activated by injecting a 0.5 mM NiCl₂ solution in the running buffer. The IKKα-His protein, at a concentration of 20 ng/µL in running buffer, was injected at a flow rate of 10 µL/min with a contact time of 60 seconds. The 5’ Biotin A+B RNA was similarly diluted in running buffer prior to injection. To ensure stable responses, three start-up cycles were performed over both the control and active surfaces before the analysis started. For regeneration, a solution of running buffer containing 0.35 M EDTA was employed to remove the nickel and any chelated molecules from the chip surface. For the inverse analysis, the 5’ Biotin A+B RNA was captured on an NA (NeutrAvidin) chip (Cytiva, cat. No. 29407997) following the manufacture instructions and [56], and varying doses of IKKα-His protein, diluted in the running buffer, were injected over the captured RNA. Sensorgrams were globally fit to 1:1 binding model using Biacore T-200 evaluation software version 1.0[57]. The injections were performed three times independently for each interaction and the mean and standard deviation of ka, kd and KD calculated.

### Purification of full-length recombinant NEMO, IKKα protein and the coiled-coil domains of IKK complex

The purification of GST-NEMO was performed as described previously[58]. In brief, the pGEX-NEMO plasmid was transformed into Rosetta BL21 cells, which were cultured in LB medium containing 100 μg/ml ampicillin. NEMO protein expression was induced by adding 0.3 mM IPTG at 28°C for 3 hours, and the protein was purified using glutathione agarose beads.

For the purification of full-length IKKα protein, the IKKα cDNA was first subcloned into the pMAL-c5X vector using mutagenesis. After sequence confirmation of the correct clone, the pMAL-c5X-IKKα-6His construct was transformed into Shuffle T7 competent E. coli cells (NEB, cat. No.C3026J), and the cells were cultured in LB medium with 100 μg/ml ampicillin. Protein expression was induced with 0.1 mM IPTG at 28°C for 16-18 hours. IKKα protein was purified from inclusion bodies. Cells were harvested by centrifugation at 4000 RPM for 20 minutes at 4°C and lysed in 2X PBS supplemented with 1 mM DTT, 10% glycerol, and 0.1% NP-40, followed by sonication. The lysate was then centrifuged at 10,000 g for 15 minutes to remove the supernatant. The pellet was resuspended in 2X PBS containing 10% glycerol, 0.5 mM DTT, and 1% (w/v) sarkosyl, and incubated at 22°C on a rocker at 120 rpm for 16 hours, followed by sonication. The supernatant was collected and dialyzed against 2 liters of dialysis buffer (2X PBS with 0.2 mM DTT and 15 mM imidazole) to remove sarkosyl, with buffer changes twice for 16-18 hours each at 22°C. After dialysis, the supernatant was subjected to nickel affinity purification as previously described[59]. The coiled-coil domain of IKK were also subcloned into the MBP tagged plasmid backbone with different tags, respectively. And the coiled-coil domain proteins were purified by using the same method, specific to each tag.

### Bioinformatics analysis

Raw RNA-Seq files for prostate and lung cancer cell lines from the Cancer Cell Line Encyclopedia (CCLE) database were downloaded from the NCBI Bioproject PRJNA523380 [60, 61]using the prefetch tool (https://github.com/ncbi/sra-tools) and converted to FASTQ format using fasterq-dump. Quality control was performed by using the BBduk from the BBTools package (https://doi.org:https://jgi.doe.gov/data-and-tools/software-tools/bbtools/bb-tools-user-guide/) to remove adapters, and trim bases below 20 and discard reads shorter. The GRCh38 reference genome was indexed using hisat2-build[62] and aligned to reference genome using hisat2. SAM files were converted to BAM format, sorted and indexed using samtools for downstream analyses [63].

The controlled RNA-Seq BAM files for the lung adenocarcinoma from the TCGA (TCGA-LUAD) were downloaded from the Database of Genotypes and Phenotypes (dbGAP) using the gdc-client tool.

To detect DRAIC isoforms (exon 3-4a-4 and exon 3-4), specific 40-base probes were designed. For exon 3-4a-4, the probe included the last 5 bases of exon 3, 30 bases from exon 4a, and the first 5 bases of exon 4. For exon 3-4, the probe comprised the last 20 bases of exon 3 and the first 20 bases of exon 4 (sequences in Supp. Table 2). Probes were mapped to RNA-Seq reads, and isoform-specific expression was quantified as counts per million (CPM = (Raw Counts/Total Mapped Reads) × 10^6). The scaling factor 10^6 is the normalizing factor for variations in sequencing depth between samples.

Normalized RNA-Seq data (DESeq2) from TCGA-LUAD were stratified by isoform expression into low (Q1) and high (Q4) groups for exon 3-4a-4 and exon 3-4. Differential expression between groups was analyzed using GSEA v4.3.3 [63]. We used the MSigDB gene set collection, including the BIOCARTA_NFKB_PATHWAY, GOBP_NON_CANONICAL_NF_KAPPAB_SIGNAL_TRANSDUCTION, and HALLMARK_TNFA_SIGNALING_VIA_NFKB gene sets. Parameters included a maximum gene set size of 5000, minimum size of 10, 1000 permutations, and weighted enrichment statistics.

DU145 (GSM618472-GSM618477) and LNCaP (GSM618479-GSM618485) BAM files were downloaded from GEO (GSE25183) and processed according to the submitter’s instructions. Differentially expressed genes (DEGs) were visualized using the R package Pheatmap.

### Statistical analysis

All data are presented as mean ± SEM. Statistical analyses were conducted using GraphPad Prism (version 8.0.2, GraphPad Software Inc., San Diego, CA). Comparisons between two groups were performed using one-tailed, unpaired Student’s *t*-tests. For comparisons involving more than two groups, one-way or two-way ANOVA followed by Tukey’s post-hoc test was applied. A *p* value of less than 0.05 was considered statistically significant.

## Data Availability Statement

The original contributions of this study are available in the article and Supplementary Material. The SHAPE-MaP libraries data files have been deposited in the Gene Expression Omnibus (GEO) database under accession number GSE279192. For further inquiries, please contact the corresponding author.

## Author Contributions

XH, RKP and YC designed the experiments, DS performed the bioinformatics analyses and CW performed the SHAPE-Map analysis. XH, RKP, YC, AM, CW and IN performed the experiments and analyzed the data. AD and KMW supervised the study. AD and XH wrote the manuscript, and all authors reviewed and approved the final version of the manuscript.

## Acknowledgment

We express our gratitude to members of the Dutta laboratory. Special thanks to Dr. Edlue M. Tawengwa for assistance in the SPR experiment. The figures are formatted in Biorender.com. This work was supported by grants from the NIH (R01 GM146756 to A.D., and R35 GM122532 to K.M.W.), an American Cancer Society Postdoctoral Fellowship (ACS 130845-RSG-17-114-01-RMC to C.A.W.), a Predoctoral Fellowship from the American Heart Association (18PRE33990261 to R.K.P.), Wagner fellowships from the University of Virginia (to R.K.P.) and the F99/K00 NCI Predoctoral to Postdoctoral Fellow Transition Award (F99/K00CA253732 to R.K.P.). UAB Multidisciplinary Molecular Interaction Core (MMIC) is funded by NIH award 1S10RR026935

