## Supplemental Figure Legend for "A 36-base hairpin within lncRNA *DRAIC*, which is modulated by alternative splicing, interacts with the IKKα coiled-coil domain and inhibits NF-κB and tumor cell phenotypes"

**SUPPLEMENTARY FIGURE LEGENDS**

**Fig. S1** A+B specifically binds to IKKα.

**A** EMSA examines if A+B binds to proteins other than IKKα by mixing 10-fold excess of indicated other proteins with A+B RNAs.

**B** EMSA tests whether IKKα binds to another RNA with a stem-loop, but a different sequence: M3 RNA (sequence provided in the Supplemental Table1).

**Fig. S2** A+B inhibits the phosphorylation of IκBα in HeLa and C4-2B cell lines.

**A and B** Western blotting was performed to detect the phosphorylation levels of IkBα after overexpressing DRAIC derivatives in HeLa cells (A) and C4-2B (B). Ponceau S. staining shows that the equal amount of protein was loaded. The ratio of p-IκBα to total IκBα, after normalization to the ratio in the EV lanes is displayed below the blot.

**Fig. S3** Overexpressing different portions of DRAIC in PC3M cell confirmed by RT-qPCR.

RT-qPCR measures the level of DRAIC after overexpressing different portions of DRAIC in PC3M cells. Primer sequences provided in the Supplemental Table1.

**Fig. S4** A+B inhibits migration in HeLa and C4-2B cancer cells.

**A and C** Matrigel Invasion assay was conducted in HeLa (A) and C4-2B (C) cells after transfection with FL DRAIC and DRAIC derivatives to see if A+B affects tumor cell invasion. The invaded cells were quantified. Data are expressed as mean ± SD; ***P* < 0.01, *****P* < 0.0001 by one-way ANOVA followed with Tukey correction. ns= no significant when compared to EV.

**B and D** RT-qPCR measures the level of DRAIC after overexpressing different portions of DRAIC in HeLa (B) and C4-2B (D) cancer cells.

**Fig. S5** Heatmap of RNA seq data from additional cell lines (besides the pair shown in Fig. 8C) showing that high levels of exon 3-4a-4 DRAIC (left three cell lines) are associated with lower level of NF-κB target genes compared to low exon 3-4a-4.
