## Supplementary figures and images for "A 36-base hairpin within lncRNA *DRAIC*, which is modulated by alternative splicing, interacts with the IKKα coiled-coil domain and inhibits NF-κB and tumor cell phenotypes"

### Supplemental Fig.2

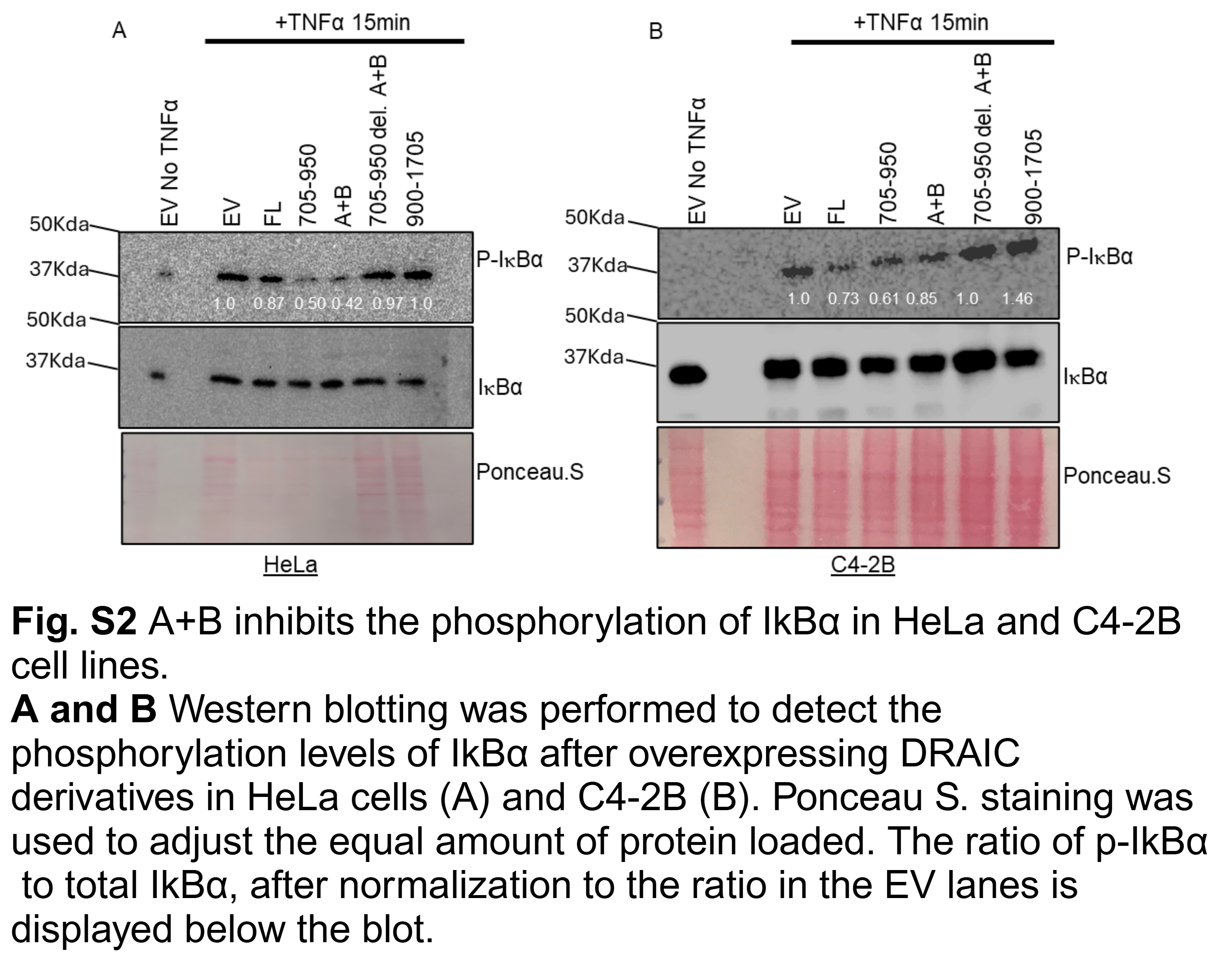

### Supplemental Fig.3

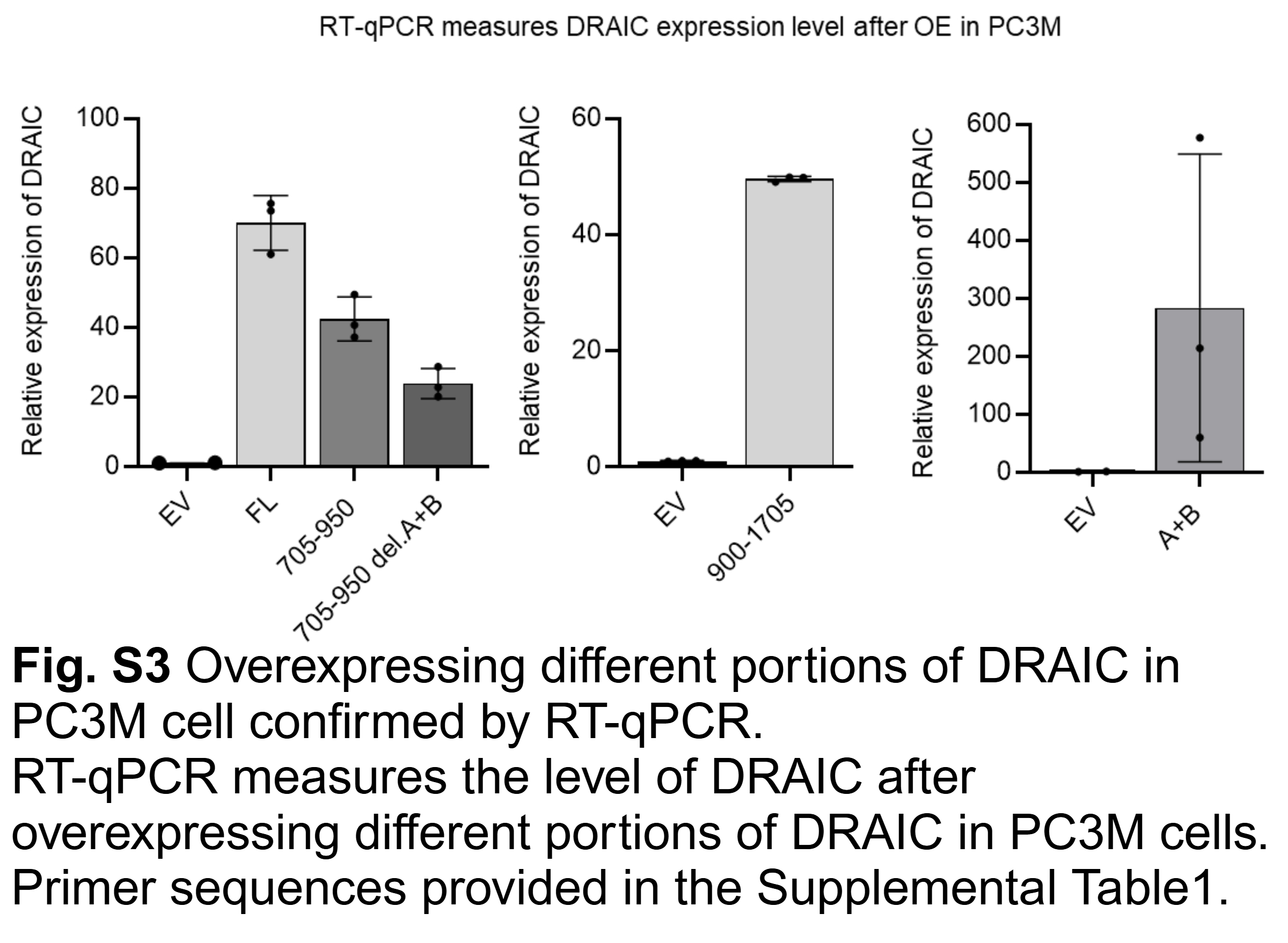

### Supplemental Fig.4

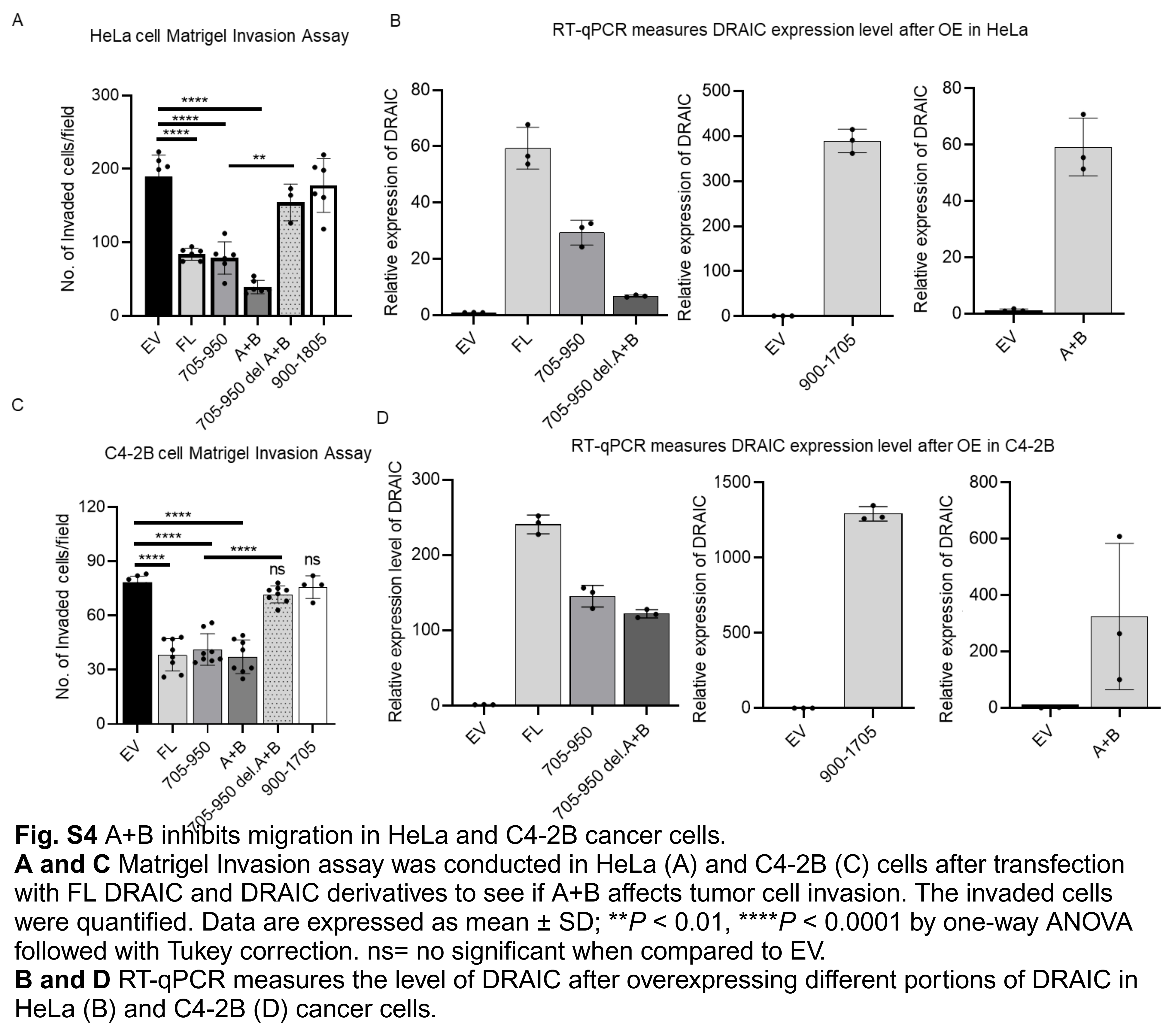

### Supplemental Fig.5

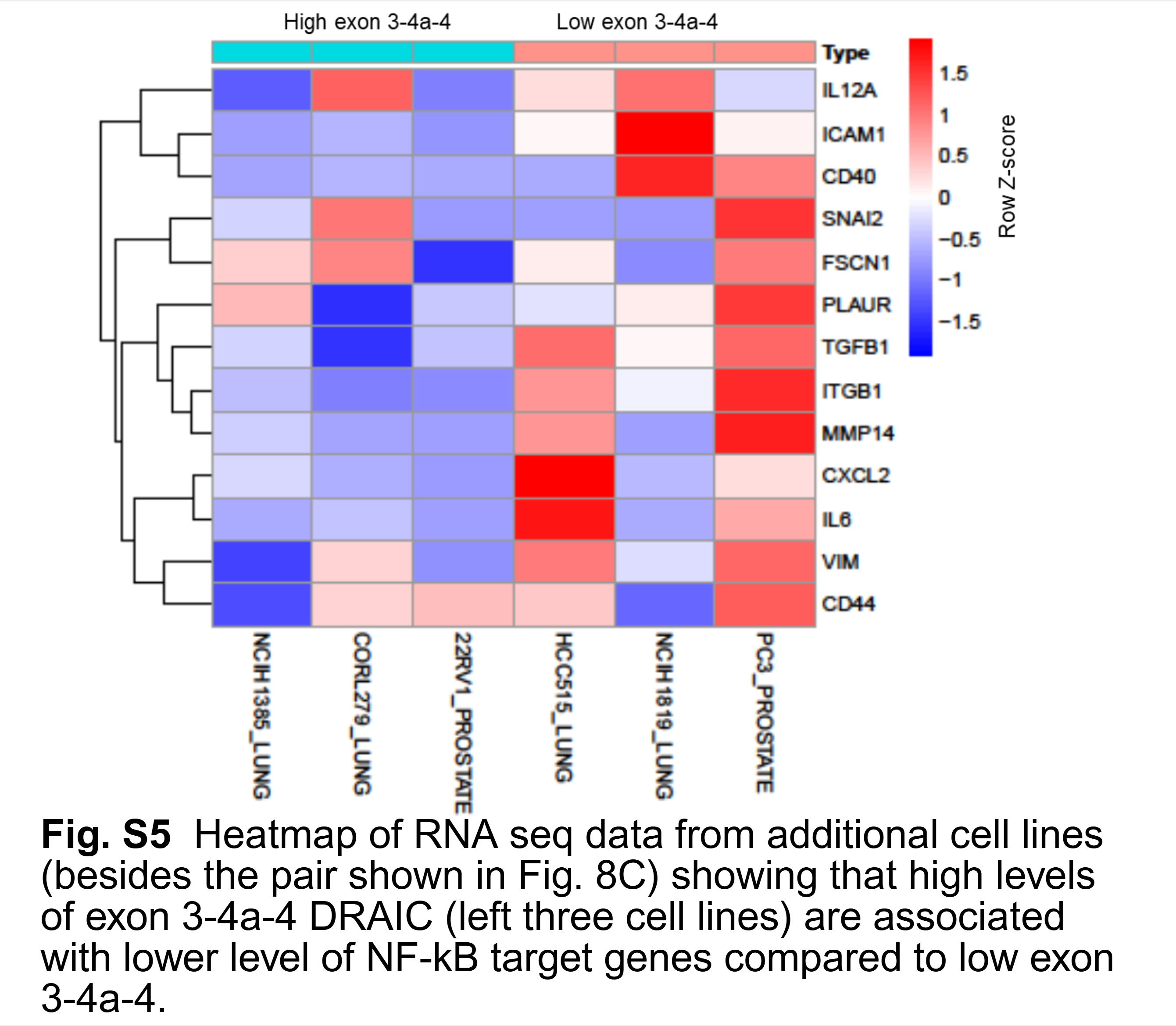

### Supplemental Figure 1

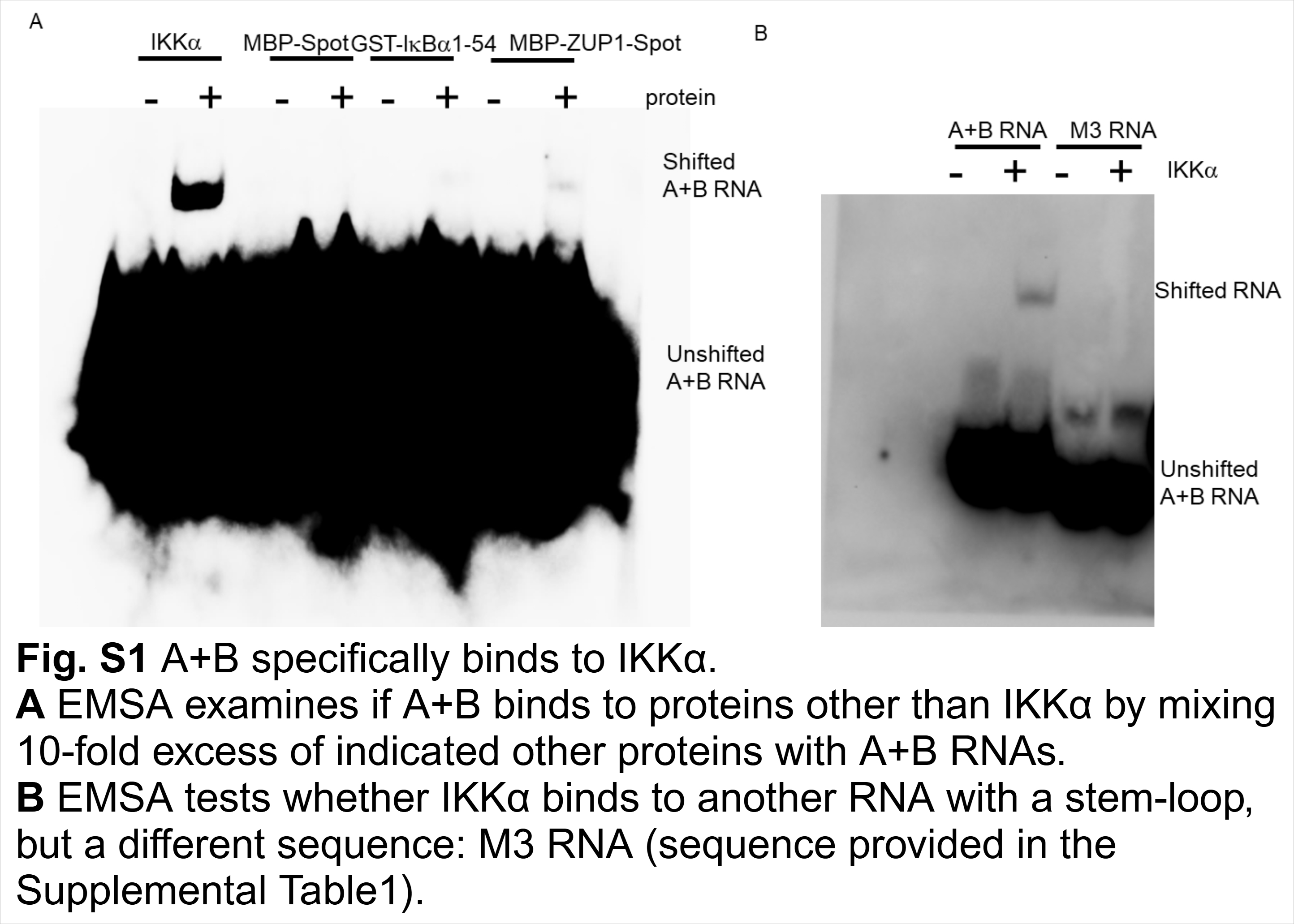
