## Supplemental Table for "A 36-base hairpin within lncRNA *DRAIC*, which is modulated by alternative splicing, interacts with the IKKα coiled-coil domain and inhibits NF-κB and tumor cell phenotypes"

**Table S1. Sequences of primers used in this study**

|  | Forward primer (5ʹ to 3ʹ) | Reverse primer (5ʹ to 3ʹ) |
| --- | --- | --- |
| APTR (qPCR) | TGGCCATTTCCACATGTCAG | CTAGGCTCTTTCTTCATC |
| APTR (IVT) | TAATACGACTCACTATAGGGAGATGGCCATTTCCACATGTCAG | GTACACTTCACACCTGAACAC |
| A+B (qPCR) | TACCGAGCTCGGATC | ATAGGGCCCTCTAGATG |
| DRAIC  (800-905) | CGCTCAACAACGAAAAGCAAAAAG | TTCATTAATGACACCTGATTTTGGAGGC |
| DRAIC  (IVT 705-950) | TAATACGACTCACTATAGGGAGATTGGAATAACTTGCAG | CATTTCTTCAGAATTTTC |
| DRAIC  (960-1105) | ATTTCCTGCTTTTCACAGAAAAC | ATGTTCATACTTCTGCTGCGTC |
| DRAIC  (IVT FL) | TAATACGACTCACTATAGGGAGACAGGGAGCTGGTTCCAGG | GACAGAGACGGAGAGATTTATTTT |
| GAPDH | AGGTCGGTGTGAACGGATTTG | TGTAGACCATGTAGTTGAGGTCA |
| Luciferase | ATGGAAGACGCCAAAAAC | CCTTATGCAGTTGCTCTCC |
| Luciferase (IVT) | TAATACGACTCACTATAGGGAGAATGGAAGATGCCAAAAACATTAAG | TTACACGGCGATCTTGCCGC |
| EV RNA sequence | GAGACCCAAGCUUGGUACCGAGCUCGGAUCCUCGAGCAUGCAUCUAGAGG | |
| A+B RNA sequence | UUGGAAUAACUUGCAGUGCCAGCAACUUGUUCACAA | |
| M3 RNA sequence | CCUUGGUGGCCCAUAGUGCCAAUGGCCCGCCAAAGG | |

**Table S2:** Sequences that are used to probe each exon.

| **exons** | **Probe sequence** |
| --- | --- |
| Exon 3-4a-4 | CTGAGGAACTTGGAATAACTTGCAGTGTCTTGCAGTATTG |
| Exon 3-4 | ATGTGTAAACACTTGCTGAGTATTGTGAAACCAGCAACTT |
